# Single-cell atlas of bronchoalveolar lavage from preschool cystic fibrosis reveals new cell phenotypes

**DOI:** 10.1101/2022.06.17.496207

**Authors:** Jovana Maksimovic, Shivanthan Shanthikumar, George Howitt, Peter F Hickey, William Ho, Casey Anttila, Daniel V. Brown, Anne Senabouth, Dominik Kaczorowski, Daniela Amann-Zalcenstein, Joseph E. Powell, Sarath C. Ranganathan, Alicia Oshlack, Melanie R. Neeland

## Abstract

Inflammation is a key driver of cystic fibrosis (CF) lung disease, not addressed by current standard care. Improved understanding of the mechanisms leading to aberrant inflammation may assist the development of effective anti-inflammatory therapy. Single-cell RNA sequencing (scRNA-seq) allows profiling of cell composition and function at previously unprecedented resolution. Herein, we seek to use multimodal single-cell analysis to comprehensively define immune cell phenotypes, proportions and functional characteristics in preschool children with CF. We analyzed 42,658 cells from bronchoalveolar lavage of 11 preschool children with CF and a healthy control using scRNA-seq and parallel assessment of 154 cell surface proteins. Validation of cell types identified by scRNA-seq was achieved by assessment of samples by spectral flow cytometry. Analysis of transcriptome expression and cell surface protein expression, combined with functional pathway analysis, revealed 41 immune and epithelial cell populations in BAL. Spectral flow cytometry analysis of over 256,000 cells from a subset of the same patients revealed high correlation in major cell type proportions across the two technologies. Macrophages consisted of 13 functionally distinct sub populations, including previously undescribed populations enriched for markers of vesicle production and regulatory/repair functions. Other novel cell populations included CD4 T cells expressing inflammatory IFNα/β and NFκB signalling genes. Our work provides a comprehensive cellular analysis of the pediatric lower airway in preschool children with CF, reveals novel cell types and provides a reference for investigation of inflammation in early life CF.

## INTRODUCTION

Inflammation is a key hallmark of lung disease in cystic fibrosis (CF). A proinflammatory state is present in early life and associated with the development of irreversible structural lung disease (1). Current treatments do not directly target pulmonary inflammation in CF, and the search for effective anti-inflammatory treatment is ongoing. A better understanding of the mechanisms underlying the aberrant inflammation seen in CF would help to develop novel approaches for anti-inflammatory therapy. This could, in turn, improve outcomes for people with CF even in the era of highly effective modulator therapy (2).

Single-cell technologies offer the opportunity to better understand cell composition and function using small volume samples. Such technologies include single-cell RNA sequencing (scRNA-seq), multiomic single-cell sequencing (including cellular indexing of transcriptomes and epitopes by sequencing (CITE-seq)), and spectral flow cytometry (an advance on conventional flow cytometry that allows analysis across the whole fluorescence spectrum). The use of scRNA-seq in CF was recently reviewed by Januska and Walsh (3) who found that the majority of studies had recruited adult participants. The authors highlighted that it was *“crucial to include infant and young children in future investigations”* (3). Further, they reported that most studies had focused on the respiratory epithelium, with only two studies using sample types primarily comprised of immune cells. Schupp *et al*, compared sputa collected from adults with CF and healthy controls (4). They found sputa from those with CF had an increased proportion of recruited monocytes and neutrophils, with a corresponding reduction in alveolar macrophages. The authors also reported functional differences in cell types in people with CF, namely immature and proinflammatory neutrophils as well as monocytes demonstrating increased heat shock responses. Li *et al* compared bronchoalveolar lavage (BAL) from adults with a mild CF lung disease to healthy controls (5). While they described a novel alveolar macrophage “supercluster” based on expression of the genes *IFI27* and *APOC2*, no differences in gene expression were observed between those with CF and controls.

To improve our understanding of pulmonary inflammation in early life CF lung disease, and to address the knowledge gaps identified by Januska and Walsh (3), we aimed to apply novel multimodal single-cell analyses to BAL samples collected from children with CF as part of clinical care in the first 6 years of life. We also included one healthy control for comparison. We applied scRNA-seq, CITE-seq and spectral flow cytometry. The results offer several advancements in our understanding of lung immune cell phenotype and composition in children with CF.

## MATERIALS AND METHODS

### Study participants

All subjects (n=11 with CF and n=1 healthy control) are enrolled in the AREST CF cohort at the Royal Children’s Hospital, Melbourne, Australia. All families gave written and informed consent for their involvement in the AREST CF research program (HREC #25054), which includes collection of samples and clinical data. The healthy control participant had no history of lower airway disease and underwent bronchoscopy to investigate upper airway pathology. Supplementary Table E1 describes the demographics of study participants, and Figure 1A outlines the experimental design.

**Figure 1.**
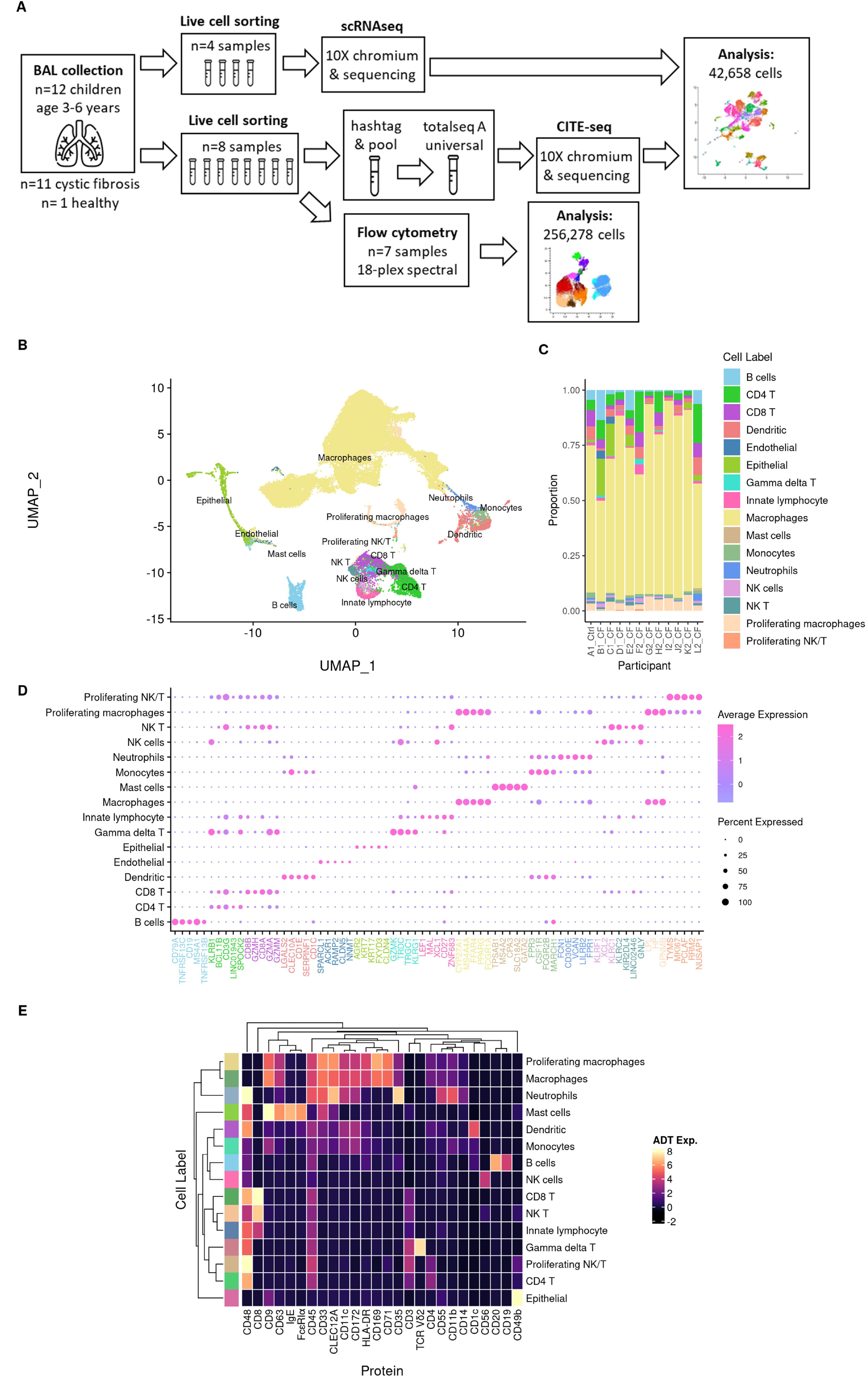
Broad immune and epithelial cell profile of the pediatric lung. **(A)** Multimodal experimental design: 12 cryopreserved bronchoalveolar (BAL) samples from children aged 3-6 years were thawed and sorted by FACS for live, single cells. These cells underwent scRNA-seq, CITE-seq, or flow cytometry as specified and were analysed to create a cell atlas of the pediatric lower airway. **(B)** UMAP visualisation of broad immune and epithelial cell populations, coloured by cell type. **(C)** Proportions of each broad population in each participant **(D)** Most significant marker genes for each broad population. **(E)** Expression of TotalSeq antibody derived tags (ADT) for immune cell lineage markers in each broad cell population for 8 samples that were analysed by CITE-seq. Endothelial cells were only identified in 2 participants whose samples were analysed by scRNA-seq, so there is no ADT data for this cell type.

### BAL sample collection and cryopreservation

All subjects underwent clinically indicated bronchoscopy. BAL was performed under general anaesthesia. Each BAL aliquot consisted of 1mL/kg (maximum 20mL) of normal saline being inserted via the working channel of the bronchoscope and then suctioned for return. The BAL samples were sent for routine bacterial culture is a clinical laboratory used to processing samples from children with CF. Viral polymerase chain reaction testing was also performed by the clinical laboratory.

The portion of the BAL samples used for research were placed on ice and processed immediately after the procedure. Samples were centrifuged at 300 x g for 10 min at 4°C. Cell-free BAL supernatant was then collected and stored at −80°C for cytokine analysis. The cell pellet was resuspended in 10mL of media (RPMI supplemented with 2% fetal calf serum (FCS)), filtered through a 70uM filter and centrifuged at 300 x g for 7mins at 4°C. Supernatant was discarded and the cell pellet resuspended in media of for cell counting. The sample was then centrifuged at 300 x g for 10 min and resuspended in equal parts media and chilled freezing solution (FCS with15% dimethyl sulfoxide (DMSO)). The freezing solution was added to resuspended cells drop by drop on ice. The samples were transferred to cryovials, and then cooled at −1°C per minute to −80 °C overnight before being transferred to liquid nitrogen for storage.

### Single-cell sequencing and flow cytometry processing

#### 1. BAL cell thawing and live single cell sorting

Cryopreserved BAL cells were thawed in 10mL media (RPMI supplemented with 10% heat-inactivated FCS) with 25U/mL benzonase at 37°C and centrifuged at 300xg for 10 min. The pellet was resuspended in 1mL PBS for cell counting. Following cell count, 9mL PBS was added to the tube and cells were centrifuged at 300 x g for 10 min. Supernatant was discarded and the cell pellet resuspended in PBS for viability staining using live/dead fixable near infrared viability dye (Invitrogen) according to manufacturers’ instructions. The viability dye reaction was stopped by the addition of FACS buffer (2% heat-inactivated FCS in PBS 2mM EDTA) and cells were centrifuged at 400 x g for 5 min. Cells were resuspended in FACS buffer for live, single cell sorting using a BD FACSAria Fusion according to the gating strategy outlined in Supplementary Figure E1.

#### 2. Sample and raw data processing for scRNA-seq (participants A1-D1)

Viable cells were sorted on a BD Influx cell sorter (Becton-Dickinson) into PBS + 0.1% bovine serum albumin and retained on ice. Sorted cells were counted and assessed for viability with Trypan Blue using a Countess automated counter (Invitrogen), and then resuspended at a concentration of 800-1000 cells/μL (8×10^5^ to 1×10^6^ cells/mL). Final cell viability estimates ranged between 92-96%. Single cell suspensions were loaded onto 10X Genomics Single Cell 3’ Chips along with the reverse transcription (RT) mastermix as per the manufacturer’s protocol for the Chromium Single Cell 3’ Library (10X Genomics; PN-120233), to generate single cell gel beads in emulsion (GEMs). Reverse transcription was performed using a C1000 Touch Thermal Cycler with a Deep Well Reaction Module (Bio-Rad) as follows: 55°C for 2h; 85°C for 5min; hold 4°C. cDNA was recovered and purified with DynaBeads MyOne Silane Beads (Thermo Fisher Scientific; Cat# 37002D) and SPRIselect beads (Beckman Coulter; Cat# B23318). Purified cDNA was amplified as follows: 98°C for 3min; 12x (98°C for 15s, 67°C for 20s, 72°C for 60s); 72°C for 60s; hold 4°C. Amplified cDNA was purified using SPRIselect beads and sheared to approximately 200bp with a Covaris S2 instrument (Covaris) using the manufacturer’s recommended parameters. Sequencing libraries were generated with unique sample indices (SI) for each chromium reaction. Libraries for all samples were multiplexed and sequenced across on 2×150 cycle flow cells on an Illumina NovaSeq 6000 (26bp (Read 1), 8bp (Index), and 98 bp (Read 2)).

The Cell Ranger Single Cell Software Suite (version 6.0.2) by 10x Genomics was used to process raw sequence data into FASTQ files. First, raw base calls from multiple flow cells were demultiplexed into separate pools of samples. Reads from each pool were then mapped to the GRCh38/hg38 genome (version 12) using STAR. The count data was processed and analyzed in *R* as described below. This dataset will be henceforth referred to as the “scRNA-seq data”.

#### 3. Sample and raw data processing for CITE-seq (participants E2-L2)

Sorted live single cells were centrifuged at 400 x g for 5min at 4°C and resuspended in 25μL of cell staining buffer (BioLegend). Human TruStain FcX FC blocking reagent (BioLegend) was added according to manufacturers’ instructions for 10min on ice. Each tube was made up to 100μL with cell staining buffer and TotalSeq Hashtag (HTO 1-8) reagents (BioLegend) were added to each sample for 20min on ice. Cells were washed with 3mL cell staining buffer and centrifuged at 400xg for 5min at 4°C. Supernatant was discarded and each sample resuspended at 62,500 cells/100μL following which 100μL of each sample were pooled into one tube. Pooled cells were centrifuged at 400xg for 5min at 4°C, supernatant discarded, and resuspended in 25μL cell staining buffer and 25ul of TotalSeqA Human Universal Cocktail v1.0 (BioLegend) for 30min on ice. This cocktail contains 154 immune related surface proteins. Cells were washed in 3mL cell staining buffer and centrifuged at 400xg for 5min at 4°C. Following two more washes, cells were resuspended in PBS + 0.04% BSA for Chromium captures.

Single-cell captures and library preparations were processed with the 10x Genomics Chromium single-cell Platform using the 10x Chromium Next GEM single-cell 3’ Reagent V3.1 Dual Index kits (10x Genomics, USA) following the manufacturer’s manual. In brief, pooled cells were counted and an estimated 40,000 cells were loaded per lane onto the 10x Chromium controller to form single-cell Gel Beads-in-Emulsion (GEMs) in duplicate. Captured cells were then lysed and barcoded and mRNA molecules were reverse transcribed to generate cDNA within the single GEMs. The barcoded cDNA was PCR-amplified and scRNA-seq libraries were constructed using the 10x 3’v3.1 library kits. The TotalSeq-A scADT-seq and scHTO-seq libraries were constructed as per manufacturer’s instructions (BioLegend). The duplicate scRNA-seq, scADT-seq and scHTO-seq libraries were quantified by Tapestation 4200 D1000 chip (Agilent). Upon input normalisation, the 6 libraries were pooled and sequenced on the Illumina NextSeq500 sequencing platform to generate 400 million 2×75-bp paired-end reads.

Reads from each sample were processed using 10x Genomics Cell Ranger software (version 5.0.0). ‘cellranger mkfastq’ was used to demultiplex the Illumina sequencer’s BCL files into FASTQ files. Next, ‘cellranger count’ was used to generate single-cell gene-count matrices against the 10x Genomics pre-built GRCh38 reference genome and transcriptome (2020-A (July 7, 2020) version). All subsequent analysis was performed in R (version 4.0.3) (R Core Team. R. A language and environment for statistical computing, 2020) with Bioconductor (version 3.12) (6). The DropletUtils package (version 1.10.3) (7) was used to create a SingleCellExperiment object from the Cell Ranger output directories and to identify non-empty droplets. Samples were demultiplexed based on the hashtag oligo (HTO) counts using the ‘hashedDrops’ function from DropletUtils with the default parameters. This was performed separately for each capture. In parallel, samples were demultiplexed using their genetic data by running cellsnp-lite (v1.2.0) (8) and Vireo (v0.5.6) (9). Specifically, reads from non-empty droplets were genotyped at 36.6M SNPs with minor allele frequency (MAF) > 0.0005 in the 1000 Genomes Project (http://ufpr.dl.sourceforge.net/project/cellsnp/SNPlist/genome1K.phase3.SNP_AF5e4.chr1toX.hg38.vcf.gz) using cellsnp-lite and then vireo assigned each cell barcode to 1 of 8 donors, doublets, or unassigned based on these genotypes. This was performed separately for each capture and then pairs of donors were matched across captures by identifying the best match between captures based on the genotype profile of each donor. As more cells were confidently assigned to donors using genetic demultiplexing, the genetic assignments were used for downstream analysis. Almost all genetic assignments corresponded to a single HTO, which was then used to link each cell’s genetic assignment to a study participant (Supplementary Table E2). This dataset will be henceforth referred to as the “CITE-seq data”.

#### 4. Flow cytometry protocol and analysis (participants F2-L2)

BAL samples from seven participants had remaining cells for flow cytometry analysis. Following viability stain as described above, cells were then resuspended in human FC-block according to manufacturers’ instructions for 5 minutes at room temperature. The antibody cocktail (Supplementary Table E3) made up at 2X concentration was added 1:1 with the cells and incubated for 30 minutes on ice. Following staining, cells were washed with 2 mL FACS buffer and centrifuged at 400 x g for 5min. Cells were then resuspended in 2% PFA for a 20min fixation on ice, washed, and resuspended in 150μl FACS buffer for acquisition using a 5L Cytek Aurora. Our protocols for the collection and processing of pediatric BAL for single-cell analysis are publicly available at https://www.protocols.io/workspaces/earlyAIR. Flow cytometry results were analysed (manual gating, UMAP) using FlowJo Version 10.8.1 software. Manual gating was performed according to the gating strategy depicted in Supplementary Figure E1. UMAP analysis was conducted using a concatenated file containing all live single cells (total 256, 278) from each individual using default parameters within FlowJo. Mean expression levels of each protein were exported from FlowJo for each manually gated cell type, following which heatmap plotting and unsupervised hierarchical clustering was performed using the Morpheus heatmap tool (https://software.broadinstitute.org/Morpheus).

### Single-cell sequencing data analysis

The “scRNA-seq data” was generated from 4 individuals’ samples (<u>participants A1-D1</u>), across 4 captures (1 control sample, 3 CF samples). The “CITE-seq data” was generated from a further 8 individuals’ samples (<u>participants E2-L2</u>) multiplexed across 2 captures (8 CF samples). Both datasets were similarly processed using the R statistical programming language (version 4.1.0). As previously described for the “CITE-seq data” in Methods Section 2 of *Single-cell sequencing and flow cytometry processing*, the DropletUtils package (version 1.14.1) (10) was used to identify non-empty droplets in the “scRNA-seq data”. This was performed separately for each capture. There were 33,538 non-empty droplets across the 4 samples for the “scRNA-seq data” and 36,601 across the 8 samples for the “CITE-seq data”. To detect within-sample doublets that cannot be identified from the genetic and HTO demultiplexing data, additional doublets were called for each capture using both scds (version 1.10.0) (10) and scDblFinder (version 1.8.0) (11). scds was run in cxds, bcds and hybrid mode with the *estDbl* parameter set to TRUE, scDblFinder was run without cluster mode using the doublet rate calculation as implemented in Demuxafy (12). Quality control was performed on each sample independently by visually examining the total cell number, the total unique molecular identifier (UMI) count distributions, the number of unique genes detected, and the proportions of ribosomal and mitochondrial gene counts per cell. Outlier droplets in terms of mitochondrial read % were detected based on being more than 3 median absolute deviations (MADs) from the median value of the metric across all droplets, in each sample. Droplets that were identified as mitochondrial read % outliers were then removed. DropletQC (13) was used to detect and tag additional empty droplets; there was convincing evidence of additional empty droplets present in the “scRNA-seq data”. The filtered droplets from the “scRNA-seq data” and “CITE-seq data” were then combined across 32,732 unique genes, resulting in a total of 54,106 unique droplets. 1,558 “uninformative” genes, such as mitochondrial genes, ribosomal genes, sex chromosome genes and pseudogenes, were then removed. Droplets that were unassigned or heterogenic doublets, along with HTO cross-sample doublets and the additional DropletQC empty droplets originating from the “scRNA-seq data”, leaving 45,590 droplets. 19,120 genes were retained for downstream analysis after genes expressed in less than 20 cells were discarded.

The droplets were then automatically annotated with cell type labels from the Human Lung Cell Atlas (HLCA) versions 1 (https://app.azimuth.hubmapconsortium.org/app/human-lung) and 2 (https://app.azimuth.hubmapconsortium.org/app/human-lung-v2) using versions 0.4.1 and 0.4.4 of the Azimuth online application, respectively. The annotated cells were then split into 3 separate data subsets based on their HLCA level 3 labels: “macrophage” (33,161 cells), “T/NK” (6,462 cells) and all “other” cells (5,922 cells), to facilitate confirmation of cell identity and identification of cellular subpopulations. Genes without associated Entrez identifiers were removed at this stage, leaving 16,001 genes for subsequent analyses.

Each data subset (“macrophage”, “T/NK” and “other” cells) was independently normalised with SCTransform (14), integrated, scaled, and clustered using Seurat (version 4.0.6) (15–18). Data integration of the 12 samples for each subset of cells was performed using reciprocal principal components analysis (RPCA) with the 4 “scRNA-seq data” samples set as references, 30 dimensions, 3000 features and 20 neighbours for anchor selection. For the “other” cells subset, the number of neighbours to consider when weighting anchors (k.weight) was set to the smallest number of cells in a single sample minus one. The cells in each subset were clustered using the smart local moving (SLM) algorithm with 30 principal components. Ten resolutions between 0.1 and 1 were explored using the clustree (19) package (version 0.4.4); a resolution of 1 was selected for downstream analysis of each data subset. This strategy resulted in 23, 18, and 23 subclusters in the “macrophage”, “T/NK” cell, and “other” cell data subsets, respectively. The clusters were visualised using Uniform Manifold Approximation and Projection (UMAP). The quality of each cluster was assessed by examining their Azimuth prediction score distributions, total UMI count distributions and the distributions of the total number of unique genes detected.

Marker gene analysis was performed for each cluster, in each of the data subsets, as previously described in Sim et al. 2021 (20), except that a 1.5-fold change cut-off was used for the TREAT test. Gene set enrichment analysis of the REACTOME gene sets (https://www.gsea-msigdb.org/gsea/msigdb/collections.jsp) was performed for each cluster in each data subset using ‘camera’ (21). The antibody derived tag (ADT) data associated with the “CITE-seq data” was normalised using the dsb (22) method (version 1.0.1) following their suggested workflow (https://cran.r-project.org/web/packages/dsb/vignettes/end_to_end_workflow.html). The combination of the top marker genes, REACTOME pathways and ADT expression for each cluster was used to assign them cellular and/or functional identities. The clusters were then consolidated using these manually assigned labels. Cells in clusters with no obvious cellular and/or functional identity and poor-quality control metrics were removed from subsequent analyses. A “macrophage” subset cluster displaying marker genes for both macrophage and T-cell lineages was found to be overrepresented for within-sample doublets called by *both* scds and scDblFinder (Fisher’s Exact Test p-value < 0.05). All the cells in this cluster were removed, along with all other “macrophage” subset doublets called by *both* scds and scDblFinder. The remaining “macrophage” subset cells were then re-clustered and manually annotated as previously described. This strategy resulted in 12, 12 and 15 manually annotated clusters in the “macrophage”, “T/NK” and “other” subsets, respectively. Broad cell type labels were also manually assigned to each cluster, resulting in a total of 13 broad cell type labels across the three data subsets. Marker genes for the consolidated subpopulations in each data subset were identified using the ‘Cepo’ (23) method.

The “macrophage”, “T/NK” and “other” cell data subsets were subsequently combined resulting in a total of 42,658 cells. The data were then normalised, integrated, scaled and clustered as previously described herein. The subpopulation and broad cell type labels were visualised using UMAP. Marker genes for the broad cell types were identified using the ‘Cepo’ method. Cell type proportions estimated by flow cytometry and scRNA-seq were compared using the ‘propeller’ approach; the proportions were arcsin transformed and linear models fitted using limma, taking the paired individual samples into account. The Benjamini-Hochberg procedure (24) was used to adjust for multiple testing.

All of the code, figures and outputs for the analyses described herein can be viewed at the following workflowr (25) (version 1.7.0) analysis website on GitHub: https://oshlacklab.com/paed-cf-cite-seq/. The code, as well as all necessary inputs and outputs can be cloned from the GitHub repository associated with the analysis website: https://github.com/Oshlack/paed-cf-cite-seq.

## RESULTS AND DISCUSSION

### 1. Broad immune and epithelial cell profile of BAL

Unsupervised clustering of single-cell sequencing data and cell annotation using marker gene analysis revealed the following transcriptionally distinct broad cell populations in pediatric BAL: macrophages, proliferating macrophages, monocytes, dendritic cells, neutrophils, B cells, CD4 T cells, CD8 T cells, NK cells, NK-T cells, proliferating NK/T cells, innate lymphocytes, γδ T cells, mast cells, epithelial cells and endothelial cells (Figure 1B). Macrophages were the most abundant cell type identified, followed by CD4 T cells, proliferating macrophages, CD8 T cells, dendritic cells, innate lymphocytes, B cells, monocytes, NK cells and neutrophils. Epithelial cells, NK-T cells, γδ T cells, proliferating NK/T cells, mast cells, and endothelial cells were rarer, and not detected in all individuals (11/12, 11/12, 9/12, 10/12, 6/12, and 2/12 respectively) (Figure 1C).

The five most significant marker genes for each broad cell population are shown in Figure 1D. These marker genes included those described previously for respective cell populations; macrophages (CYP27A1, MS4A4A and PPARG), proliferating macrophages and proliferating NK/T cells expressing cyclic genes (PCLAF and MKI67), B cells (CD19, MS4A1), CD4 T cells (CD3D, CD3E), CD8 T cells (CD8A, CD8B), innate lymphocytes (LEF1, XCL1), γδ T cells (TRGC1, GZMK), NK cells (KLRC1, XCL1), NK-T cells (KLRC1, CD3D), dendritic cells (CLEC10A, CD1E), monocytes (CSF1R, FCGR2B), mast cells (TPSAB1, CPA3), epithelial cells (AGR2, KRT17), and endothelial cells (SPARCL1, ACKR1) (26–30).

Annotation of each cell population was further confirmed by analysis of expression of TotalSeq-A ADTs (Figure 1E). Macrophages expressed myeloid lineage proteins CD172α, CD11c, CD71, HLA-DR, and CD169 as previously reported for alveolar macrophages in adults (31, 32). Monocytes were distinguished from macrophages based on the pan monocyte marker CD14, as well as the lack of CD71 and CD169. Dendritic cells expressed CD11c, CI)172α and CD1c; and mast cells expressed known protein markers FcεR1α, CD63, and IgE (33). Cells annotated as lymphoid lineage by marker genes expressed expected lineage proteins, including B cells (CD19^+^CD20^+^), CD4 T cells (CD3^+^CD5^+^CD4^+^), CD8 T cells (CD3^+^CD5^+^CD8^+^), γδ T cells (CD3^+^TCRVδ2^+^), NK cells (CD3^-^CD56^+^) and NK-T cells (CD3^+^CD56^+^). Epithelial cells were positive for tissue resident marker CD49b. Endothelial cells were not identified by ADTs as they were only detected in transcriptomic data from two participants, neither of whom were included in the CITE-seq experiment (Figure 1C).

### 2. Comparison of cell proportions by single-cell sequencing and flow cytometry

To investigate how results obtained by single-cell sequencing compared to those obtained by flow cytometry, we directly compared the 7 BAL samples analysed with both techniques. Our flow cytometry panel permitted identification of macrophages, monocytes, dendritic cells, neutrophils, B cells, CD4 T cells, CD8 T cells and airway epithelial cells across 256,000 live, single cells (Figure 2A, Supplementary Figure E1). Proportions of macrophages, monocytes, B cells, CD4 T cells, CD8 T cells and airway epithelial cells were generally consistent between the two techniques for each individual (Figure 2B-C). However, we observed statistically significant differences in the proportion of neutrophils (fold change flow vs scRNA-seq: 1.2, FDR=0.006) and dendritic cells (fold change flow vs scRNA-seq: −1.17, FDR=0.019) (Figure 2C). Due to the low RNA content of neutrophils and previously reported challenges in characterising these cells using single-cell technologies such as 10x (34), the higher proportion of neutrophils detected by flow cytometry was unsurprising. The reduced proportion of dendritic cells detected by flow cytometry compared to scRNA-seq was likely due to the absence of key dendritic cell subset markers CD1c and CD123 in our flow cytometry panel. Comparable protein expression patterns were observed for common surface markers CD3, CD8, CD4, CD19, CD16, CD45, HLADR, CD11c, CD14, CD47 and CD279 across the two techniques (Figure 2D). Of note, our flow cytometry panel also included several surface proteins not available in the TotalSeq-A cocktail, including CD206 for macrophages, CD15 and CD66b for neutrophil subtyping, and EPCAM for airway epithelial cells (Figure 2C). Airway macrophages showed auto-fluorescent signatures in flow cytometry data as we and others have described previously (35, 36).

**Figure 2.**
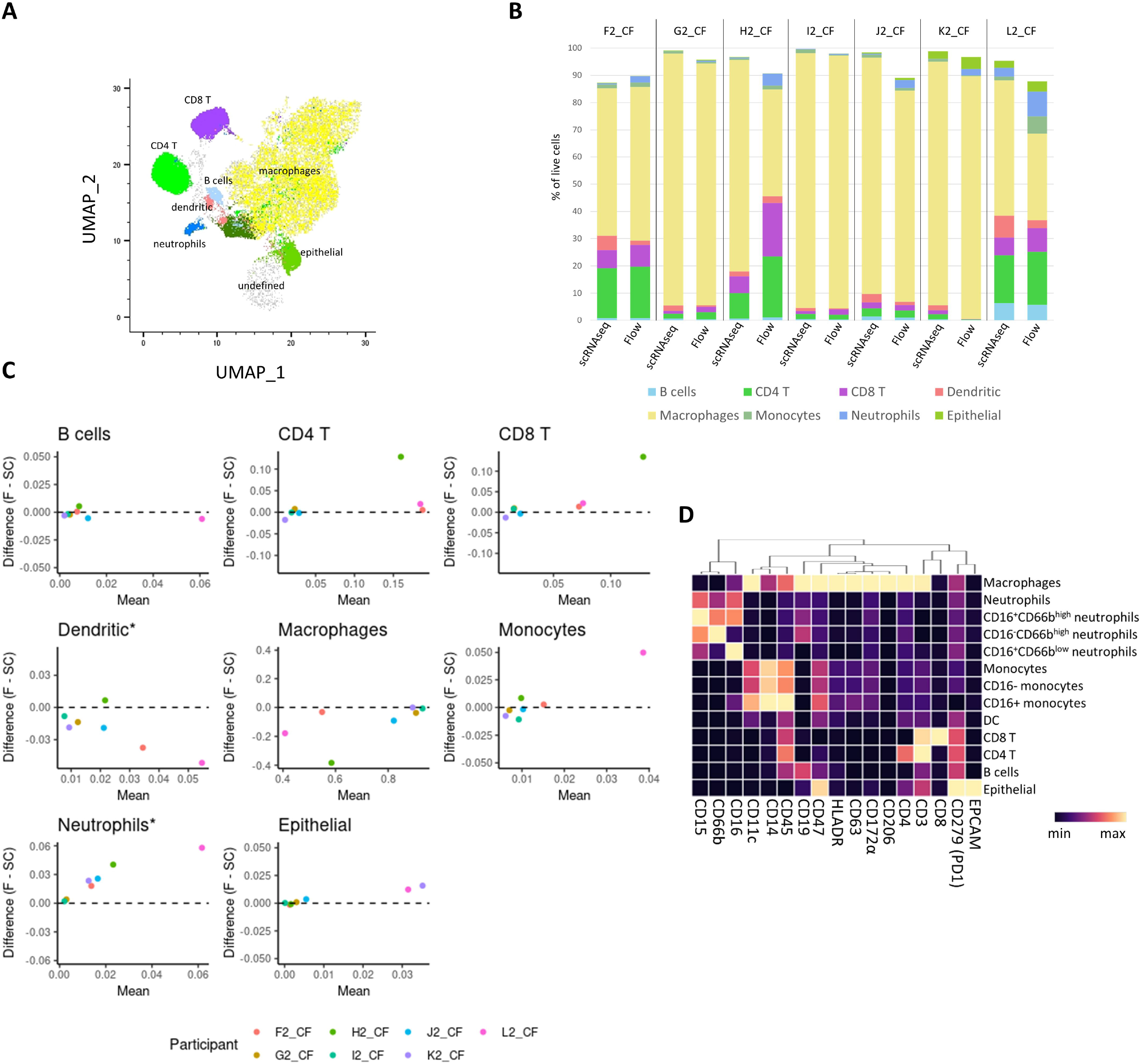
Flow cytometry analysis of BAL samples and comparison with scRNA-seq. **(A)** UMAP depicting major cell populations identified by spectral flow cytometry of 7 samples: B cells, CD4 T cells, CD8 T cells, dendritic cells, macrophages, monocytes, neutrophils and airway epithelial cells. **(B)** Cell proportions identified by scRNA-seq and flow cytometry in samples from individuals that received both techniques (participants F2-L2). Common cell types identified in both datasets are shown. All cell types identified by scRNA-seq are shown in Figure 1. **(C)** Statistical comparison of the difference in proportions identified by flow cytometry and scRNA-seq. *indicates cell types where a significant difference was observed. **(D)** Cell type specific protein expression by flow cytometry for markers included in the panel: CD15, CD66b, CD16, CD11c, CD14, CD45, CD19, CD47, HLA-DR, CD63, CD172α, CD206, CD4, CD4, CD8, CD279 (PD1), and EPCAM.

### 3. Identification of distinct lung macrophage subpopulations

As macrophages are the most abundant immune cell in BAL and are known to undergo significant functional development in early life (37), we performed a subclustering analysis of our broad macrophage population (Figure 3A-B). Subclusters were annotated based on marker gene analysis (Figure 3C), ADT protein expression (Figure 3D), and with reference to what has been described in previous scRNA-seq literature. To further confirm annotations and understand subpopulation functionality based on cell population marker genes, we also performed REACTOME pathway enrichment analyses for each subtype (38). A list of enriched REACTOME pathways in each population is provided in Supplementary Table E4, and a full list of cluster marker genes can be found in extended data file 1.

**Figure 3.**
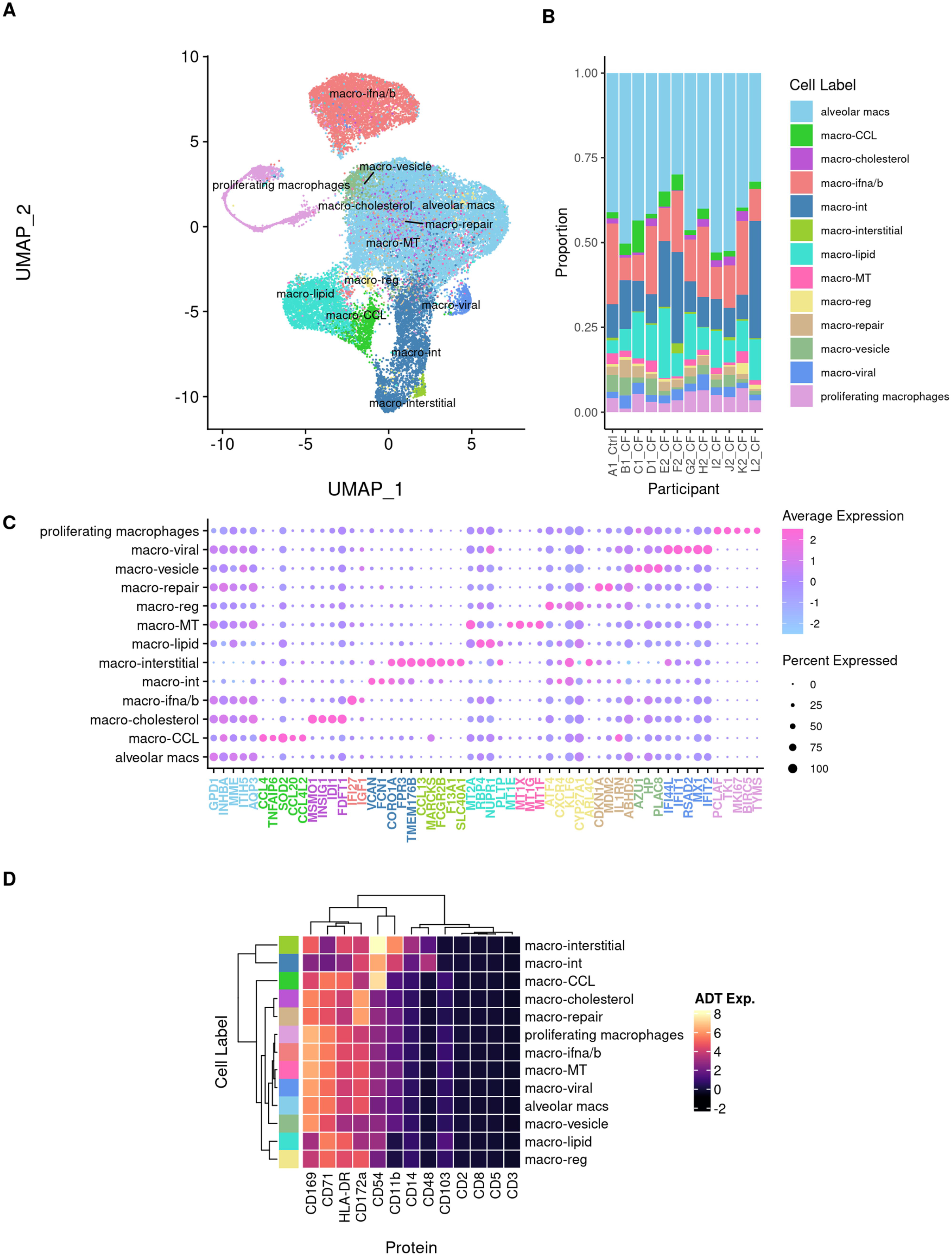
Identification of functionally distinct macrophage subpopulations. **(A)** UMAP visualisation of macrophage subpopulations identified by a subclustering analysis of cells within the total macrophage pool. **(B)** Proportions (as a percent of all macrophages) of each macrophage subpopulation in each participant. **(C)** Most significant marker genes for each macrophage subpopulation. **(D)** Expression of relevant TotalSeq antibody derived tags (ADT) in each macrophage subpopulation for 8 samples that were analysed by CITE-seq.

Our analysis revealed 13 distinct subpopulations of macrophages, of which several have recently been described in adults. All macrophage subclusters were identified in all participants including the healthy control (Figure 3B). Subtypes characterised in the adult lung and also identified here include intermediate macrophages expressing both monocyte and macrophage genes (macro-int) (27), chemokine-expressing macrophages (macro-CCL) (27), interstitial macrophages (26), proliferating (or cycling) macrophages (5), macrophages associated with lipid digestion and transport (NUPR1, RBP4) denoted as macro-lipid (39), macrophages expressing cholesterol biosynthesis related genes (MSM01, FDFT1) denoted as macro-cholesterol (5), macrophages expressing IFN-α/β signalling genes (macro-IFN-α/β) (5), macrophages expressing genes associated with viral response (macro-viral)(5) and metallothionein-expressing macrophages (macro-MT) (27) (Figure 3B). Novel subtypes identified were macrophages expressing genes associated with; vesicle production (AZU1, PLAC8) denoted as macro-vesicle; regulatory function (ATF4, GDF15) denoted as macro-reg; and response to DNA damage (CDKN1A, MDM2) denoted macro-repair (Figure 3C). Interestingly, macro-interstitial, macro-CCL, and macro-int cells expressed high protein levels of CD54 (ICAM1), a transmembrane glycoprotein receptor known to play an important role in migration of leukocytes to sites of inflammation (40) (Figure 3D).

The most abundant macrophage subcluster in our data, at a median of 16.1% of macrophages, was a population expressing IFN-α/β signalling genes that we denoted as macro-IFN-α/β. This cluster was characterised by expression of IFI27 and IGF1 and was enriched for REACTOME pathways including interferon-alpha-beta signalling. It has previously been identified in analysis of BAL from adults with CF (5).

The macro-lipid (9.4% of macrophage) subcluster was defined based on enrichment of genes (NUPR1, RBP4) and REACTOME pathways associated with lipid digestion and transport. Based on the results of gene set enrichment analysis, this cluster appears to be similar to one identified previously. Further, RBP4 was identified as a marker gene in both the current and previous study (39). Similarly, the macro-cholesterol subpopulation appears similar to a previously reported subcluster in terms of gene set enrichment analysis and a shared marker gene (MSM01)(5). The macro-viral subcluster was a rare population, which was also identified in a prior study of BAL from adults from CF where it was referred to as IFN-reacting (5).

The macro-int population was previously reported in a study of BAL from adults with COVID-19, where those with more severe disease had an increased proportion of this subtype (39). In our study this population accounted for 11.2% of the macrophages. In addition to expressing genes associated with both monocytes and macrophages, the macro-int population also expressed proteins associated with both populations, including CD11b, CD48 and intermediate levels of CD14 (Figure 3D). Similarly, the macro-interstitial population also expressed CD14 and CD11b protein (Figure 3D). CD14 expressing macrophages were also identified in our flow cytometry data (Supplementary Figure E2A). In general, proportions of CD14^+^ macrophages identified by flow cytometry and sc-RNAseq were similar (Supplementary Figure E2B) although this requires further validation using markers such as CD11b, CD48, and CD54 to differentiate the macro-int from macro-interstitial using flow cytometry.

The macro-CCL population in our dataset was enriched for the chemokine-receptor-bind-chemokines REACTOME pathway, and expressed genes for chemokines CCL4, CCL20, CCL23, CXCL5, CXCL8, CXCL9 and CXCL10, further confirming its annotation. This subcluster has been reported in all prior BAL based scRNA-seq studies, however we did not identify the 4 macro-CCL subclusters that were reported by Li *et al* (5).

The macro-MT population has been previously identified by scRNA-seq evaluation of BAL samples (5, 41). It was proposed that the gene expression profile of this subpopulation indicates these cells have been exposed to heavy metals (such as zinc, cadmium, or iron) (41). If this is correct, the presence of this population in preschool children indicates that this exposure, and subsequent development of the subpopulation of macrophages, occurs in early life.

Overall, our data suggest macrophages in the preschool CF lung are highly activated. This is evidenced by the identified subclusters, but also enrichment for a range of inflammatory pathways including IL-1 signalling, IFNα/β activation, chemokine signalling, and antiviral responses. This is consistent with prior studies showing that the upper airway of children demonstrates a baseline pre-activated state, characterised by upregulation of antiviral and inflammatory signatures compared to adults (42). Our findings demonstrate that children with CF may have these pre-activated signatures in the lower airway as well.

### 4. Neutrophil-like population identified by scRNA-seq

Given the importance of neutrophilic inflammation in CF we attempted to identify neutrophils in the single-cell data (1). Despite the aforementioned challenges of studying neutrophils by scRNA-seq, we identified a population of 252 neutrophil-like cells, to which 7 donors contributed more than 10 cells and the remaining 5 fewer than 10 cells. This contrasts with the only study which has used scRNA-seq to profile BAL from people with CF, which did not identify a neutrophil population (5). The population of cells we identified mapped to neutrophils identified by a publicly available and annotated scRNA-seq dataset of the adult lung (43) and expressed inflammatory genes including those encoding for S100 proteins and the IL-1 signalling pathway. Protein expression on these cells included CD35, CD55, CLEC12A, CD48, and a lack of HLA-DR (Figure 1E). CD55 and CD35 are known to be involved in neutrophil phagocytosis (44). Cells in this neutrophil-like population also shared gene and protein features with monocytes (genes: FCN1, CD300E; proteins: CD11b and CD14), which has been described for neutrophils previously (45–48). A more extensive panel of neutrophil-specific ADTs that are not currently included in the TotalSeq A cocktail could be used to further characterise this cell type; for example, CD15 and CD66b are protein markers that we used to distinguish BAL neutrophils from monocytes by flow cytometry (36).

We show by flow cytometry that BAL neutrophils can be further subtyped based on CD16 and CD66b expression, with CD16^-^CD66b^high^, CD16^+^CD66b^high^ and CD16^+^CD66b^low^ subsets identified (Supplementary Figure E1, Figure E2D). This classification has previously been used to identify Granule-Releasing Immunomodulatory Metabolically active neutrophils (CD16^-^CD66B^high^) which has been implicated in CF related lung inflammation (49). These cells are characterised by primary granule release and show impaired capacity for bacterial clearance.

An inherent limitation of our data is that our samples were cryopreserved and thawed prior to analysis. This process is known to deplete granulocytes; however, we have shown in previous work that cryopreservation of BAL should not affect the yield of other immune cell populations (35). As is the case with most clinical studies, our sampling times are unpredictable in nature and cryopreservation is unavoidable. However, the use of fresh samples would not have obviated issues with identifying neutrophil populations in scRNA-seq data, as evidenced by the absence of neutrophils in the prior study of CF BAL which used fresh samples (5).

### 5. Inflammatory CD4 T cell subsets in the pediatric lower airway

Our initial gene and protein expression analysis in Figure 1 showed broad populations of CD4 T cells, CD8 T cells, innate lymphocytes, NK cells, NK-T cells and γδ T cells in BAL of preschool children with CF. As with macrophages, we next explored heterogeneity within these T/NK subsets using a subclustering approach (Figure 4). Our analysis revealed both known and novel subtypes, using marker genes (Figure 4C, extended data file 2), ADT protein expression (Figure 4D) and REACTOME pathway analysis (Supplementary Table E5). Previously reported subtypes observed by single-cell RNA sequencing of adult lung samples that we also observed in our analysis were CD8 Trm, CD8-GZMK, CD4 Treg, γδ T cells, NK cells, NK-T cells, innate lymphocytes and proliferating NK/T cells (26, 27, 43).

**Figure 4.**
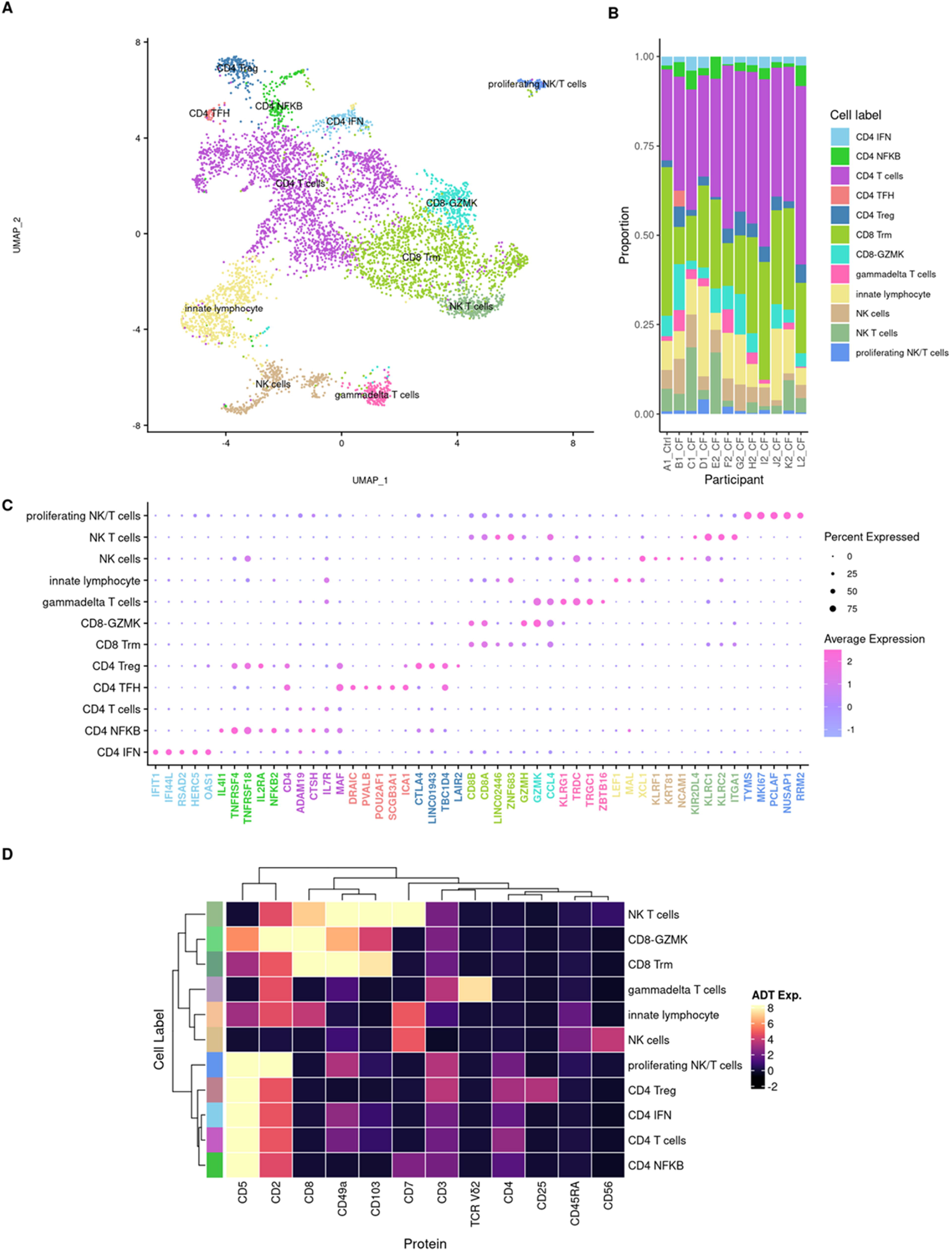
Characterisation of distinct T and NK cell subsets. **(A)** UMAP visualisation of T and NK cell subsets identified by a subclustering analysis of cells within the ‘T/NK’ group. **(B)** Proportions (as a percent of all T/NK cells) of each T/NK subpopulation in each participant. **(C)** Most significant marker genes for each identified T and NK cell subset. **(D)** Expression of TotalSeq antibody derived tags (ADT) for T and NK cell markers in each cell subset for 8 samples that were analysed by CITE-seq. CD4 TFH cells were only identified in 1 participant whose sample was analysed by scRNA-seq, so there is no ADT data for this cell type.

Two subtypes of CD8 T cells (CD8 Trm and CD8-GZMK) and NK-T cells showed gene and protein markers of lung tissue residency, including surface expression of CD103 and CD49a (50) (Figure 4C-D). Conversely, CD4 T cell subsets, NK cells, innate lymphocytes, and γδ T cells were negative or only weakly positive for CD103 and CD49a, suggesting they may have been recently recruited from the circulation. Both CD8 Trm and CD8-GZMK subtypes expressed genes encoding for cytotoxic proteases GZMA, GZMB, GZMH, GZMM however CD8-GZMK was the only CD8 subtype to express GZMK. Recent work has shown that patterns of cytotoxic molecule expression relate to CD8 T cell differentiation stage, with a lack of GZM expression observed in naïve CD8 T cells, the majority of memory CD8 T cells co-expressing GZMA/B/M, and a low number of intermediately-differentiated CD8 T cells expressing GZMK (51). CD4 T follicular helper (CD4-TFH) cells expressing TFH genes SCGB3A1 and PD1 signalling (52) were also identified, however only in one individual (Figure 4B).

To our knowledge, we are the first to describe subtypes of CD4 T cells enriched for IFN and NFκB signalling genes in the lung. Features of the CD4 T-IFN subtype include marker genes IFIT1, IFIT3, RSAD2 (Figure 4C) as well as REACTOME enrichment for interferon and antiviral pathways. Features of the CD4 T-NFκB subtype include marker genes TNFRSF4, TNFRSF18, NFKB2 (Figure 4C) as well as REACTOME enrichment for several NFκB activation and signalling pathways (Supplementary Table E5). These inflammatory CD4 T cells were observed in both the healthy control and children with CF (Figure 4B).

This highlights that these cells may represent a unique characteristic of the early life CF lung and adds further evidence to the concept that the pediatric CF lower airway is primed for inflammatory responses.

### 6. Defining rare cells

A final subclustering analysis of all other cells (not macrophages, not T/NK) revealed further heterogeneity within the rarer DC, B cell and epithelial cell clusters (Figure 5, Supplementary Table E6, extended data file 3). DCs were separated into conventional DC (cDC1 and cDC2), plasmacytoid DC, and migratory DC, based on previously described marker genes (26) and protein expression. cDC1 expressed CLEC9A gene and CD141 protein, whilst cDC2 expressed CD1c (53–55). Plasmacytoid DC expressed plasmacytoid surface marker CD123, were negative for conventional markers CD1c and CD141, and marker genes included CLEC4C and GZMB. Plasma B cells were also identified, characterised by key marker genes (JCHAIN, MZB1, TNFRSF17) as well as protein expression of plasma markers CD27 and CD38 (56) (Figure 5C-D).

**Figure 5.**
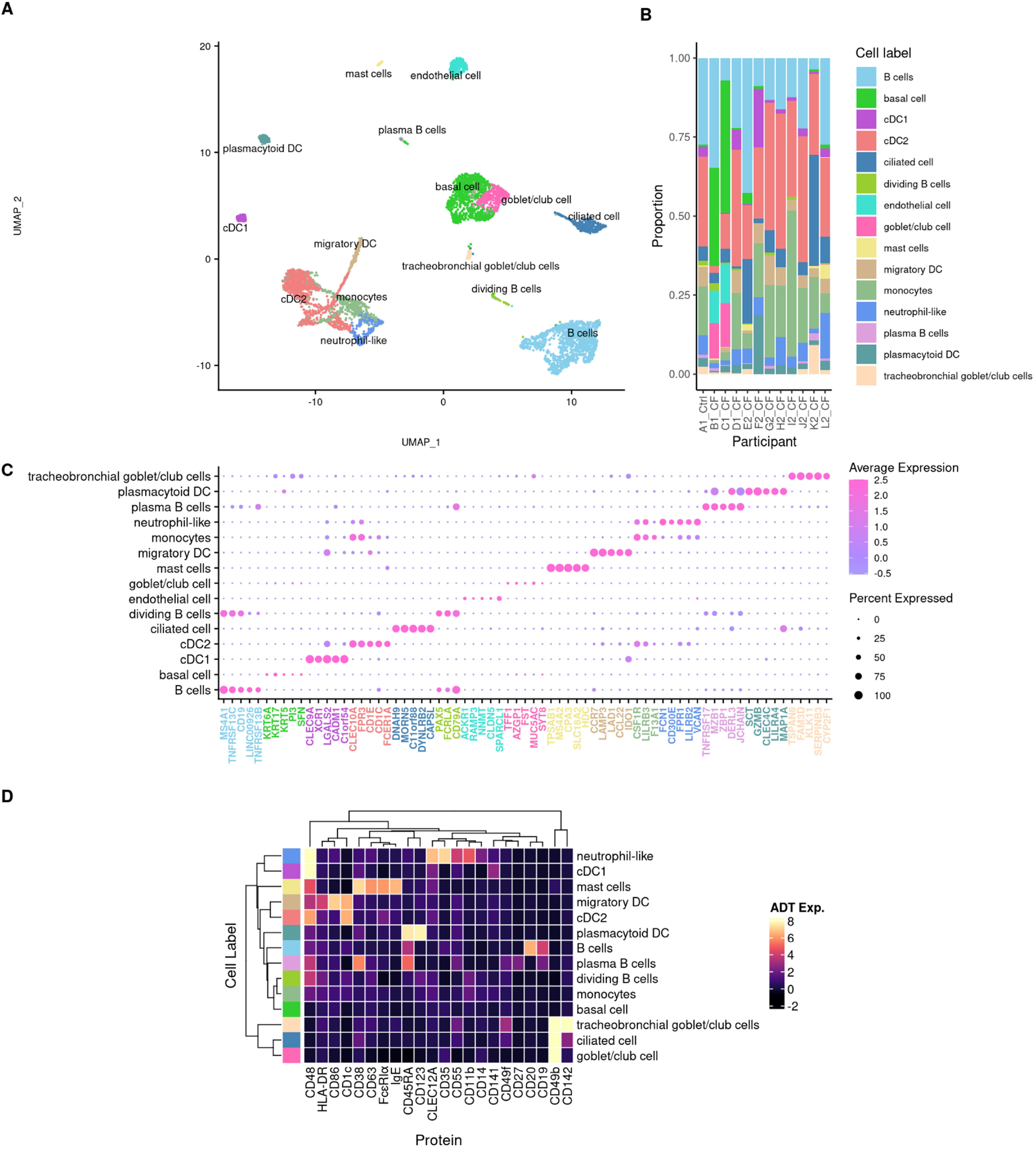
Characterisation of rare myeloid, B cell and epithelial cell population. **(A)** UMAP visualisation of cell subsets identified by a subclustering analysis of cells within the ‘not macrophage, not T/NK’ population. **(B)** Proportions (as a percent of all “other” cells) of each “other” subpopulation in each participant. **(C)** Most significant marker genes for each cell subset. **(D)** Expression of relevant TotalSeq antibody derived tags (ADT) in each cell subset for 8 samples that were analysed by CITE-seq. Endothelial cells were only identified in 2 participants whose samples were analysed by scRNA-seq, so there is no ADT data for this cell type.

Whilst BAL predominantly samples immune cells of the lower airway, a small fraction of airway epithelial cells was also captured. We identified 4 subtypes of airway epithelial cells: ciliated epithelial cells, basal epithelial cells and 2 clusters containing cells of a goblet/club phenotype. These were annotated using the Human Lung Cell Atlas (HCLA) (26). The ciliated epithelial cell cluster expressed marker genes which are exclusively expressed by ciliated columnar cells of the tracheobronchial tree, and multiciliated epithelial cells in the HLCA. The basal epithelial cells expressed well-validated marker genes KRT17, KRT5, and KRT6a. Two distinct populations expressing a mixture of goblet and club cell marker genes were identified. One cluster, denoted as goblet/club cells, expressed marker genes including MUC5AC, TFF1, AZGP1, FST, and SYT8 which are expressed by club, goblet and mucus secreting cells. Another cluster, denoted as tracheobronchial goblet/club cells, expressed the marker gene TSPAN8 which is a HCLA marker gene for both club and goblet cells from the lower airway.

## CONCLUSIONS

This is the first single-cell analysis of bronchoalveolar lavage from preschool children with CF and includes both RNA and protein. These novel data fill a substantial gap in existing pulmonary scRNA-seq data and in CF lung disease.

Early life is a crucial period of development for both epithelial and immune cells. For example, unlike other organs such as the skin, liver and brain where the transcriptome of resident macrophages from fetal and adult life are very similar, there are substantial differences in the gene expression profiles of fetal and adult lung resident macrophages (57). This highlights childhood as a period where fetal lung resident macrophages must develop before eventually exhibiting an adult phenotype.

Previous scRNA-seq studies involving people with CF have characterised adult samples and primarily focused on respiratory epithelium rather than immune cells. This is also true for the broader pulmonary scRNA-seq field (3). For example, the recent release of the HLCA contained a core dataset derived from samples of 107 individuals, however the youngest of these individuals was 10 years old (26). The HLCA also included an extended dataset, however only 6 of the 338 (1.7%) samples were from individuals 6 years or younger and these samples were not analysed separately to identify specific developmental changes.

Integration of transcriptome-wide data, assessment of highly multiplexed surface proteins, and functional pathway analysis, enabled us to extensively characterise 41 cell populations in the bronchoalveolar lavage of preschool children, predominantly with CF. We revealed several novel cell subtypes, most notably functional macrophage subpopulations and inflammatory CD4 T cells. These novel cell types were found in both children with CF and the one control participant, implying they are unlikely to be disease specific, although this needs to be confirmed in larger studies. Our analysis of paired cell surface protein expression using TotalSeq ADTs and spectral flow cytometry further extends current knowledge of lung cell types and provides within-study validation of our transcriptomic findings.

There are several future directions resulting from this work. First, replication in a larger cohort of children with CF and control participants will allow both validation of our findings as well as investigation of differences in health and disease. In addition, given the variability of the CF lung disease phenotype, it may be possible to identify transcriptomic signatures associated with disease severity. It will also be important to further validate the novel cell populations identified in our study in other cohorts and by different technologies.

This study provides the first transcriptional profile of early life lower pulmonary samples and provides a reference dataset for researchers investigating the pulmonary immune system in early life cystic fibrosis lung disease.

## ACKNOWLEDGEMENTS

The authors would like to acknowledge the generous support and guidance provided by Professor Daniel Chambers and Dr Brendan O’Sullivan regarding experimental methods when designing the first study. We thank the children and parents who participated in the AREST-CF study, without whom this work would not have been possible. Finally, we thank the Chan Zuckerberg Initiative Single-cell Biology community for helpful discussions and support.

## REFERENCES

1. Sly PD, Gangell CL, Chen L, Ware RS, Ranganathan S, Mott LS, Murray CP, Stick SM. Risk factors for bronchiectasis in children with cystic fibrosis. The New England journal of medicine 2013; 368: 1963–1970.

2. Shanthikumar S, Ranganathan S, Neeland MR. Ivacaftor, not ivacaftor/lumacaftor, associated with lower pulmonary inflammation in preschool cystic fibrosis. Pediatric pulmonology 2022; 57: 2549–2552.

3. Januska MN, Walsh MJ. Single-Cell RNA-Sequencing Reveals New Basic and Translational Insights in the Cystic Fibrosis Lung. American journal of respiratory cell and molecular biology 2022.

4. Schupp JC, Khanal S, Gomez JL, Sauler M, Adams TS, Chupp GL, Yan X, Poli S, Zhao Y, Montgomery RR, Rosas IO, Dela Cruz CS, Bruscia EM, Egan ME, Kaminski N, Britto CJ. Single-Cell Transcriptional Archetypes of Airway Inflammation in Cystic Fibrosis. American journal of respiratory and critical care medicine 2020; 202: 1419–1429.

5. Li X, Kolling FW, Aridgides D, Mellinger D, Ashare A, Jakubzick CV. ScRNA-seq expression of IFI27 and APOC2 identifies four alveolar macrophage superclusters in healthy BALF. Life science alliance 2022; 5.

6. Huber W, Carey VJ, Gentleman R, Anders S, Carlson M, Carvalho BS, Bravo HC, Davis S, Gatto L, Girke T, Gottardo R, Hahne F, Hansen KD, Irizarry RA, Lawrence M, Love MI, MacDonald J, Obenchain V, Oleś AK, Pagès H, Reyes A, Shannon P, Smyth GK, Tenenbaum D, Waldron L, Morgan M. Orchestrating high-throughput genomic analysis with Bioconductor. Nature methods 2015; 12: 115–121.

7. Lun ATL, Riesenfeld S, Andrews T, Dao TP, Gomes T, participants in the 1st Human Cell Atlas J, Marioni JC. EmptyDrops: distinguishing cells from empty droplets in dropletbased single-cell RNA sequencing data. Genome biology 2019; 20: 63–63.

8. Huang X, Huang Y. Cellsnp-lite: an efficient tool for genotyping single cells. Bioinformatics 2021.

9. Huang Y, McCarthy DJ, Stegle O. Vireo: Bayesian demultiplexing of pooled single-cell RNA-seq data without genotype reference. Genome Biol 2019; 20: 273.

10. Bais AS, Kostka D. scds: computational annotation of doublets in single-cell RNA sequencing data. Bioinformatics 2020; 36: 1150–1158.

11. Germain PL, Lun A, Garcia Meixide C, Macnair W, Robinson MD. Doublet identification in single-cell sequencing data using scDblFinder. F1000Res 2021; 10: 979.

12. Neavin D, Senabouth A, Hang Lee JT, Ripoll A, Franke L, Prabhakar S, Ye CJ, McCarthy DJ, Melé M, Hemberg M, Powell JE. <em>Demuxafy</em>: Improvement in droplet assignment by integrating multiple single-cell demultiplexing and doublet detection methods. bioRxiv 2022: 2022.2003.2007.483367.

13. Muskovic W, Powell JE. DropletQC: improved identification of empty droplets and damaged cells in single-cell RNA-seq data. Genome Biology 2021; 22: 329.

14. Hafemeister C, Satija R. Normalization and variance stabilization of single-cell RNA-seq data using regularized negative binomial regression. Genome Biology 2019; 20: 296.

15. Hao Y, Hao S, Andersen-Nissen E, Mauck WM, 3rd, Zheng S, Butler A, Lee MJ, Wilk AJ, Darby C, Zager M, Hoffman P, Stoeckius M, Papalexi E, Mimitou EP, Jain J, Srivastava A, Stuart T, Fleming LM, Yeung B, Rogers AJ, McElrath JM, Blish CA, Gottardo R, Smibert P, Satija R. Integrated analysis of multimodal single-cell data. Cell 2021; 184: 3573–3587.e3529.

16. Butler A, Hoffman P, Smibert P, Papalexi E, Satija R. Integrating single-cell transcriptomic data across different conditions, technologies, and species. Nat Biotechnol 2018; 36: 411–420.

17. Stuart T, Butler A, Hoffman P, Hafemeister C, Papalexi E, Mauck WM, 3rd, Hao Y, Stoeckius M, Smibert P, Satija R. Comprehensive Integration of Single-Cell Data. Cell 2019; 177: 1888–1902.e1821.

18. Stuart T, Satija R. Integrative single-cell analysis. Nat Rev Genet 2019; 20: 257–272.

19. Zappia L, Oshlack A. Clustering trees: a visualization for evaluating clusterings at multiple resolutions. GigaScience 2018; 7.

20. Sim CB, Phipson B, Ziemann M, Rafehi H, Mills RJ, Watt KI, Abu-Bonsrah KD, Kalathur RKR, Voges HK, Dinh DT, Ter Huurne M, Vivien CJ, Kaspi A, Kaipananickal H, Hidalgo A, Delbridge LMD, Robker RL, Gregorevic P, Dos Remedios CG, Lal S, Piers AT, Konstantinov IE, Elliott DA, El-Osta A, Oshlack A, Hudson JE, Porrello ER. Sex-Specific Control of Human Heart Maturation by the Progesterone Receptor. Circulation 2021.

21. Wu D, Smyth GK. Camera: a competitive gene set test accounting for inter-gene correlation. Nucleic acids research 2012; 40: e133–e133.

22. Mulè MP, Martins AJ, Tsang JS. Normalizing and denoising protein expression data from droplet-based single cell profiling. Nat Commun 2022; 13: 2099.

23. Kim HJ, Wang K, Chen C, Lin Y, Tam PPL, Lin DM, Yang JYH, Yang P. Uncovering cell identity through differential stability with Cepo. Nature Computational Science 2021; 1: 784–790.

24. Benjamini Y, Hochberg Y. Controlling the False Discovery Rate - a Practical and Powerful Approach to Multiple Testing. J R Stat Soc B 1995; 57: 289–300.

25. Blischak JD, Carbonetto P, Stephens M. Creating and sharing reproducible research code the workflowr way. F1000Research 2019; 8: 1749.

26. Sikkema L, Strobl D, Zappia L, Madissoon E, Markov N, Zaragosi L, Ansari M, Arguel M, Apperloo L, Bécavin C, Berg M, Chichelnitskiy E, Chung M, Collin A, Gay A, Hooshiar Kashani B, Jain M, Kapellos T, Kole T, Mayr C, von Papen M, Peter L, Ramírez-Suástegui C, Schniering J, Taylor C, Walzthoeni T, Xu C, Bui L, de Donno C, Dony L, Guo M, Gutierrez A, Heumos L, Huang N, Ibarra I, Jackson N, Kadur Lakshminarasimha Murthy P, Lotfollahi M, Tabib T, Talavera-Lopez C, Travaglini K, Wilbrey-Clark A, Worlock K, Yoshida M, Consortium LBN, Desai T, Eickelberg O, Falk C, Kaminski N, Krasnow M, Lafyatis R, Nikolíc M, Powell J, Rajagopal J, Rozenblatt-Rosen O, Seibold M, Sheppard D, Shepherd D, Teichmann S, Tsankov A, Whitsett J, Xu Y, Banovich N, Barbry P, Duong T, Meyer K, Kropski J, Pe’er D, Schiller H, Tata P, Schultze J, Misharin A, Nawijn M, Luecken M, Theis F. An integrated cell atlas of the human lung in health and disease. bioRxiv 2022: 2022.2003.2010.483747.

27. Madissoon E, Oliver AJ, Kleshchevnikov V, Wilbrey-Clark A, Polanski K, Orsi AR, Mamanova L, Bolt L, Richoz N, Elmentaite R, Pett JP, Huang N, He P, Dabrowska M, Pritchard S, Tuck L, Prigmore E, Knights A, Oszlanczi A, Hunter A, Vieira SF, Patel M, Georgakopoulos N, Mahbubani K, Saeb-Parsy K, Clatworthy M, Bayraktar OA, Stegle O, Kumasaka N, Teichmann SA, Meyer KB. A spatial multi-omics atlas of the human lung reveals a novel immune cell survival niche. bioRxiv 2021: 2021.2011.2026.470108.

28. Liao Z, Jin Y, Chu Y, Wu H, Li X, Deng Z, Feng S, Chen N, Luo Z, Zheng X, Bao L, Xu Y, Tan H, Zhao L. Single-cell transcriptome analysis reveals aberrant stromal cells and heterogeneous endothelial cells in alcohol-induced osteonecrosis of the femoral head. Commun Biol 2022; 5: 324.

29. Crinier A, Dumas P-Y, Escalière B, Piperoglou C, Gil L, Villacreces A, Vély F, Ivanovic Z, Milpied P, Narni-Mancinelli É, Vivier É. Single-cell profiling reveals the trajectories of natural killer cell differentiation in bone marrow and a stress signature induced by acute myeloid leukemia. Cellular & molecular immunology 2021; 18: 1290–1304.

30. Mattiola I, Tomay F, De Pizzol M, Silva-Gomes R, Savino B, Gulic T, Doni A, Lonardi S, Astrid Boutet M, Nerviani A, Carriero R, Molgora M, Stravalaci M, Morone D, Shalova IN, Lee Y, Biswas SK, Mantovani G, Sironi M, Pitzalis C, Vermi W, Bottazzi B, Mantovani A, Locati M. The macrophage tetraspan MS4A4A enhances dectin-1-dependent NK cell-mediated resistance to metastasis. Nat Immunol 2019; 20: 1012–1022.

31. Bissonnette EY, Lauzon-Joset J-F, Debley JS, Ziegler SF. Cross-Talk Between Alveolar Macrophages and Lung Epithelial Cells is Essential to Maintain Lung Homeostasis. Frontiers in immunology 2020; 11.

32. Morrell ED, Wiedeman A, Long SA, Gharib SA, West TE, Skerrett SJ, Wurfel MM, Mikacenic C. Cytometry TOF identifies alveolar macrophage subtypes in acute respiratory distress syndrome. JCI Insight 2018; 3.

33. Kraft S, Jouvin M-H, Kulkarni N, Kissing S, Morgan ES, Dvorak AM, Schröder B, Saftig P, Kinet J-P. The tetraspanin CD63 is required for efficient IgE-mediated mast cell degranulation and anaphylaxis. Journal of immunology (Baltimore, Md: 1950) 2013; 191: 2871–2878.

34. Grieshaber-Bouyer R, Radtke FA, Cunin P, Stifano G, Levescot A, Vijaykumar B, Nelson-Maney N, Blaustein RB, Monach PA, Nigrovic PA, Aguilar O, Allan R, Astarita J, Austen KF, Barrett N, Baysoy A, Benoist C, Brown BD, Buechler M, Buenrostro J, Casanova MA, Chowdhary K, Colonna M, Crowl T, Deng T, Desland F, Dhainaut M, Ding J, Dominguez C, Dwyer D, Frascoli M, Gal-Oz S, Goldrath A, Johanson T, Jordan S, Kang J, Kapoor V, Kenigsberg E, Kim J, Kim Kw, Kiner E, Kronenberg M, Lanier L, Laplace C, Lareau C, Leader A, Lee J, Magen A, Maier B, Maslova A, Mathis D, McFarland A, Merad M, Meunier E, Monach PA, Mostafavi S, Muller S, Muus C, Ner-Gaon H, Nguyen Q, Novakovsky G, Nutt S, Omilusik K, Ortiz-Lopez A, Paynich M, Peng V, Potempa M, Pradhan R, Quon S, Ramirez R, Ramanan D, Randolph G, Regev A, Rose SA, Seddu K, Shay T, Shemesh A, Shyer J, Smilie C, Spidale N, Subramanian A, Sylvia K, Tellier J, Turley S, Vijaykumar B, Wagers A, Wang C, Wang PL, Wroblewska A, Yang L, Yim A, Yoshida H, ImmGen C. The neutrotime transcriptional signature defines a single continuum of neutrophils across biological compartments. Nat Commun 2021; 12: 2856.

35. Shanthikumar S, Burton M, Saffery R, Ranganathan SC, Neeland MR. Single Cell Flow Cytometry Profiling of Bronchoalveolar Lavage in Children. American journal of respiratory cell and molecular biology 2020.

36. Shanthikumar S, Ranganathan SC, Saffery R, Neeland MR. Mapping Pulmonary and Systemic Inflammation in Preschool Aged Children With Cystic Fibrosis. Frontiers in immunology 2021; 12: 733217.

37. Tan SYS, Krasnow MA. Developmental origin of lung macrophage diversity. Development 2016; 143: 1318–1327.

38. Gillespie M, Jassal B, Stephan R, Milacic M, Rothfels K, Senff-Ribeiro A, Griss J, Sevilla C, Matthews L, Gong C, Deng C, Varusai T, Ragueneau E, Haider Y, May B, Shamovsky V, Weiser J, Brunson T, Sanati N, Beckman L, Shao X, Fabregat A, Sidiropoulos K, Murillo J, Viteri G, Cook J, Shorser S, Bader G, Demir E, Sander C, Haw R, Wu G, Stein L, Hermjakob H, D’Eustachio P. The reactome pathway knowledgebase 2022. Nucleic acids research 2021; 50: D687–D692.

39. Liao M, Liu Y, Yuan J, Wen Y, Xu G, Zhao J, Cheng L, Li J, Wang X, Wang F, Liu L, Amit I, Zhang S, Zhang Z. Single-cell landscape of bronchoalveolar immune cells in patients with COVID-19. Nature medicine 2020; 26: 842–844.

40. Figenschau SL, Knutsen E, Urbarova I, Fenton C, Elston B, Perander M, Mortensen ES, Fenton KA. ICAM1 expression is induced by proinflammatory cytokines and associated with TLS formation in aggressive breast cancer subtypes. Scientific reports 2018; 8: 11720.

41. Mould KJ, Moore CM, McManus SA, McCubbrey AL, McClendon JD, Griesmer CL, Henson PM, Janssen WJ. Airspace Macrophages and Monocytes Exist in Transcriptionally Distinct Subsets in Healthy Adults. American journal of respiratory and critical care medicine 2021; 203: 946–956.

42. Loske J, Röhmel J, Lukassen S, Stricker S, Magalhães VG, Liebig J, Chua RL, Thürmann L, Messingschlager M, Seegebarth A, Timmermann B, Klages S, Ralser M, Sawitzki B, Sander LE, Corman VM, Conrad C, Laudi S, Binder M, Trump S, Eils R, Mall MA, Lehmann I. Pre-activated antiviral innate immunity in the upper airways controls early SARS-CoV-2 infection in children. Nat Biotechnol 2022; 40: 319–324.

43. Zilionis R, Engblom C, Pfirschke C, Savova V, Zemmour D, Saatcioglu HD, Krishnan I, Maroni G, Meyerovitz CV, Kerwin CM, Choi S, Richards WG, De Rienzo A, Tenen DG, Bueno R, Levantini E, Pittet MJ, Klein AM. Single-Cell Transcriptomics of Human and Mouse Lung Cancers Reveals Conserved Myeloid Populations across Individuals and Species. Immunity 2019; 50: 1317–1334.e1310.

44. Vandendriessche S, Cambier S, Proost P, Marques PE. Complement Receptors and Their Role in Leukocyte Recruitment and Phagocytosis. Frontiers in Cell and Developmental Biology 2021; 9.

45. Abdel-Salam BK, Ebaid H. Expression of CD11b and CD18 on polymorphonuclear neutrophils stimulated with interleukin-2. Cent Eur J Immunol 2014; 39: 209–215.

46. Haziot A, Tsuberi BZ, Goyert SM. Neutrophil CD14: biochemical properties and role in the secretion of tumor necrosis factor-alpha in response to lipopolysaccharide. J Immunol 1993; 150: 5556–5565.

47. Silva-Gomes R, Mapelli SN, Boutet MA, Mattiola I, Sironi M, Grizzi F, Colombo F, Supino D, Carnevale S, Pasqualini F, Stravalaci M, Porte R, Gianatti A, Pitzalis C, Locati M, Oliveira MJ, Bottazzi B, Mantovani A. Differential expression and regulation of MS4A family members in myeloid cells in physiological and pathological conditions. J Leukoc Biol 2022; 111: 817–836.

48. Munthe-Fog L, Hummelshoj T, Honoré C, Moller ME, Skjoedt MO, Palsgaard I, Borregaard N, Madsen HO, Garred P. Variation in FCN1 affects biosynthesis of ficolin-1 and is associated with outcome of systemic inflammation. Genes & Immunity 2012; 13: 515–522.

49. Margaroli C, Moncada-Giraldo D, Gulick DA, Dobosh B, Giacalone VD, Forrest OA, Sun F, Gu C, Gaggar A, Kissick H, Wu R, Gibson G, Tirouvanziam R. Transcriptional firing represses bactericidal activity in cystic fibrosis airway neutrophils. Cell reports Medicine 2021; 2: 100239.

50. Zheng MZM, Wakim LM. Tissue resident memory T cells in the respiratory tract. Mucosal Immunol 2021.

51. Bengsch B, Ohtani T, Herati RS, Bovenschen N, Chang K-M, Wherry EJ. Deep immune profiling by mass cytometry links human T and NK cell differentiation and cytotoxic molecule expression patterns. J Immunol Methods 2018; 453: 3–10.

52. Weinstein JS, Lezon-Geyda K, Maksimova Y, Craft S, Zhang Y, Su M, Schulz VP, Craft J, Gallagher PG. Global transcriptome analysis and enhancer landscape of human primary T follicular helper and T effector lymphocytes. Blood 2014; 124: 3719–3729.

53. Minoda Y, Virshup I, Leal Rojas I, Haigh O, Wong Y, Miles JJ, Wells CA, Radford KJ. Human CD141+ Dendritic Cell and CD1c+ Dendritic Cell Undergo Concordant Early Genetic Programming after Activation in Humanized Mice In Vivo. Frontiers in Immunology 2017; 8.

54. Tullett KM, Lahoud MH, Radford KJ. Harnessing Human Cross-Presenting CLEC9A+XCR1+ Dendritic Cells for Immunotherapy. Frontiers in immunology 2014; 5.

55. Collin M, McGovern N, Haniffa M. Human dendritic cell subsets. Immunology 2013; 140: 22–30.

56. da Silva FAR, Pascoal LB, Dotti I, Setsuko Ayrizono MdL, Aguilar D, Rodrigues BL, Arroyes M, Ferrer-Picon E, Milanski M, Velloso LA, Fagundes JJ, Salas A, Leal RF. Whole transcriptional analysis identifies markers of B, T and plasma cell signaling pathways in the mesenteric adipose tissue associated with Crohn’s disease. J Transl Med 2020; 18: 44.

57. Bian Z, Gong Y, Huang T, Lee CZW, Bian L, Bai Z, Shi H, Zeng Y, Liu C, He J, Zhou J, Li X, Li Z, Ni Y, Ma C, Cui L, Zhang R, Chan JKY, Ng LG, Lan Y, Ginhoux F, Liu B. Deciphering human macrophage development at single-cell resolution. Nature 2020; 582: 571–576.

