## Supplementary material for "Single-cell atlas of bronchoalveolar lavage from preschool cystic fibrosis reveals new cell phenotypes": Online Data Supplement

[1] Computational Biology Program, Peter MacCallum Cancer Centre, Parkville, VIC, Australia
[2] Respiratory Diseases, Murdoch Children's Research Institute, Parkville, VIC, Australia
[3] Department of Paediatrics, University of Melbourne, Parkville, VIC, Australia
[4] Respiratory and Sleep Medicine, Royal Children's Hospital, Parkville, VIC, Australia
[5] Cellular Genomics Projects Team, Advanced Technology and Biology Division, WEHI, Parkville, VIC, Australia
[6] Department of Medical Biology, University of Melbourne, Parkville, VIC, Australia
[7] Garvan-Weizmann Centre for Cellular Genomics, Garvan Institute of Medical Research, Sydney, NSW, Australia
[8] UNSW Cellular Genomics Futures Institute, University of New South Wales, Sydney, NSW, Australia
[9] Sir Peter MacCallum Department of Oncology, University of Melbourne, Parkville, VIC, Australia
[10] School of Mathematics and Statistics, University of Melbourne, Parkville, VIC, Australia
[11] Molecular Immunity, Murdoch Children's Research Institute, Parkville, VIC, Australia
*Contributed equally
#Corresponding author

**Corresponding author details:**
Dr Melanie Neeland, PhD
Royal Children's Hospital, 50 Flemington Road, Parkville, VIC, Australia


### SUPPLEMENTARY TABLES AND FIGURES

**Supplementary Table E1.** Study cohort and demographics

| Participant | Sex | Age (years) | CF or control | Organisms in BAL | scRNA-seq | Totalseq ADT |
|---|---|---|---|---|---|---|
| **A1_Ctrl** | M | 3.00 | Control | Staphylococcus aureus, Haemophilus haemolyticus | Y | |
| **B1_CF** | M | 2.99 | CF | Stenotrophomonas maltophilia, Enterobacter cloacae, Burkholderia gladioli, Respiratory syncytial virus, Aspergillus species | Y | |
| **C1_CF** | M | 2.99 | CF | Bordetella pertussis | Y | |
| **D1_CF** | M | 3.03 | CF | None present | Y | |
| **E2_CF** | F | 5.99 | CF | Staphylococcus aureus, Stenotrophomonas maltophilia, Escherichia coli, Haemophilus influenzae, Parainfluenzae Virus | Y | Y |
| **F2_CF** | F | 6.02 | CF | Mycobacterium Avium, Haemophilus parainfluenzae | Y | Y |
| **G2_CF** | F | 4.91 | CF | Haemophilus parainfluenzae, Aspergillus Fumigatus | Y | Y |
| **H2_CF** | F | 5.89 | CF | None present | Y | Y |
| **I2_CF** | F | 5.93 | CF | None present | Y | Y |
| **J2_CF** | M | 5.05 | CF | Staphylococcus aureus | Y | Y |
| **K2_CF** | M | 4.92 | CF | None present | Y | Y |
| **L2_CF** | M | 5.95 | CF | Staphylococcus aureus, Haemophilus parainfluenzae, Stenotrophomonas maltophilia | Y | Y |

**Supplementary Table E2.** Concordance between HTO and genetic assignments.

| HTO / genetic donor | donor_A | donor_B | donor_C | donor_D | donor_E | donor_F | donor_G | donor_H | Doublet | Unknown | Total |
|---|---|---|---|---|---|---|---|---|---|---|---|
| **HTO_1** | 0 | 1620 | 3 | 1 | 1 | 0 | 9 | 0 | 16 | 15 | 1665 |
| **HTO_2** | 0 | 2 | 1 | 1 | 1 | 0 | 2763 | 2 | 49 | 30 | 2849 |
| **HTO_3** | 1 | 1 | 1 | 0 | 3267 | 0 | 0 | 2 | 17 | 8 | 3297 |
| **HTO_4** | 1 | 0 | 1663 | 0 | 0 | 0 | 0 | 0 | 7 | 2 | 1673 |
| **HTO_5** | 0 | 0 | 1 | 2 | 2 | 2913 | 0 | 1 | 30 | 8 | 2957 |
| **HTO_6** | 0 | 0 | 1 | 1861 | 0 | 1 | 0 | 1 | 8 | 6 | 1878 |
| **HTO_7** | 0 | 1 | 0 | 1 | 0 | 0 | 1 | 2310 | 14 | 6 | 2333 |
| **HTO_8** | 2274 | 1 | 3 | 4 | 2 | 1 | 3 | 2 | 30 | 81 | 2401 |
| **Doublet** | 181 | 163 | 248 | 216 | 380 | 330 | 273 | 281 | 1581 | 29 | 3682 |
| **Unknown** | 133 | 86 | 482 | 227 | 178 | 47 | 63 | 155 | 384 | 338 | 2093 |
| **Total** | 2590 | 1874 | 2403 | 2313 | 3831 | 3292 | 3112 | 2754 | 2136 | 523 | 24828 |

**Supplementary Table E3.** Flow cytometry antibody cocktail

| Surface Marker | Fluorophore | Clone | Final Dilution |
|---|---|---|---|
| CD16 | BUV395 | 3G8 | 1:400 |
| SIRPα | BUV563 | SE5A5 | 1:100 |
| CD47 | BUV737 | B6H12 | 1:00 |
| CD206 | BV421 | G10F5 | 1:50 |
| HLADR | V500 | G46-6 | 1:100 |
| CD19 | BV605 | SJ25C1 | 1:100 |
| CD8 | BV650 | RPA-T8 | 1:100 |
| CD45 | BV711 | HI30 | 1:50 |
| CD14 | BV786 | M5E2 | 1:25 |
| EPCAM | BB515 | EBA-1 | 1:100 |
| CD3 | FITC | SK7 | 1:50 |
| PD1 | BB700 | EH12.1 | 1:100 |
| CD66b | PE | 19.2 | 1:100 |
| CD15 | PECF594 | W6D3 | 1:100 |
| CD11c | PECy7 | B-ly6 | 1:50 |
| CD63 | A647 | H5C6 | 1:50 |
| CD4 | A700 | RPA-T4 | 1:25 |

**Supplementary Table E4.** REACTOME analysis of macrophage subclusters.

| macro-IFNα/β | Direction | PValue | FDR |
|---|---|---|---|
| REACTOME_INTERFERON_ALPHA_BETA_SIGNALING | Up | 1.09E-45 | 7.36E-43 |
| REACTOME_INTERFERON_SIGNALING | Up | 1.40E-11 | 4.71E-09 |
| **macro-lipid** | | | |
| REACTOME_LIPID_DIGESTION_MOBILIZATION_AND_TRANSPORT | Up | 4.68E-26 | 1.05E-23 |
| REACTOME_HDL_MEDIATED_LIPID_TRANSPORT | Up | 1.81E-24 | 3.06E-22 |
| **macro-int** | | | |
| REACTOME_CREATION_OF_C4_AND_C2_ACTIVATORS | Down | 2.09E-51 | 7.05E-49 |
| REACTOME_INITIAL_TRIGGERING_OF_COMPLEMENT | Down | 3.74E-32 | 6.30E-30 |
| **macro-CCL** | | | |
| REACTOME_CHEMOKINE_RECEPTORS_BIND_CHEMOKINES | Up | 3.40E-94 | 2.29E-91 |
| REACTOME_PEPTIDE_LIGAND_BINDING_RECEPTORS | Up | 1.18E-31 | 3.99E-29 |
| **macro-vesicle** | | | |
| REACTOME_AMYLOIDS | Up | 4.15E-44 | 2.79E-41 |
| REACTOME_INITIAL_TRIGGERING_OF_COMPLEMENT | Up | 1.63E-35 | 5.48E-33 |
| **maco-viral** | | | |
| REACTOME_INTERFERON_ALPHA_BETA_SIGNALING | Up | 0 | 0 |
| REACTOME_INTERFERON_SIGNALING | Up | 1.65E-137 | 5.58E-135 |
| **macro-MT** | | | |
| REACTOME_INTERFERON_GAMMA_SIGNALING | Up | 4.52E-07 | 3.04E-04 |
| **macro-repair** | | | |
| REACTOME_AKT_PHOSPHORYLATES_TARGETS_IN_THE_CYTOSOL | Up | 1.63E-181 | 1.10E-178 |
| REACTOME_PIP3_ACTIVATES_AKT_SIGNALING | Up | 2.71E-77 | 9.14E-75 |
| **macro-cholesterol** | | | |
| REACTOME_CHOLESTEROL_BIOSYNTHESIS | Up | 0 | 0 |
| REACTOME_GLYCOPROTEIN_HORMONES | Up | 4.20E-25 | 1.42E-22 |
| **proliferating-macrophage** | | | |
| REACTOME_APC_C_CDH1_MEDIATED_DEGRADATION_OF_CDC20_AND_OTHER_APC_C_CDH1_TARGETED_PROTEINS_IN_LATE_MITOSIS_EARLY_G1 | Up | 9.24E-10 | 4.85E-07 |
| REACTOME_APC_C_CDC20_MEDIATED_DEGRADATION_OF_MITOTIC_PROTEINS | Up | 1.44E-09 | 4.85E-07 |
| **macro-reg** | | | |
| REACTOME_CREATION_OF_C4_AND_C2_ACTIVATORS | Down | 2.73E-188 | 1.84E-185 |
| REACTOME_INITIAL_TRIGGERING_OF_COMPLEMENT | Down | 1.04E-89 | 3.51E-87 |
| **macro-interstitial** | | | |
| REACTOME_COMMON_PATHWAY | Up | 2.06E-31 | 1.39E-28 |
| REACTOME_FORMATION_OF_FIBRIN_CLOT_CLOTTING_CASCADE | Up | 4.80E-14 | 1.62E-11 |

*the two most significant (FDR) pathways per functional subcluster are presented

**Supplementary Table E5**. REACTOME analysis of T/NK subclusters

| **CD4 T** | **Direction** | **PValue** | **FDR** |
|---|---|---|---|
| REACTOME_CHEMOKINE_RECEPTORS_BIND_CHEMOKINES | Down | 1.12E-06 | 7.56E-04 |
| REACTOME_IMMUNOREGULATORY_INTERACTIONS_BETWEEN_A_LYMPHOID_AND_A_NON_LYMPHOID_CELL | Down | 0.01666626 | 0.999955616 |
| **CD8 Trm** | | | |
| REACTOME_IMMUNOREGULATORY_INTERACTIONS_BETWEEN_A_LYMPHOID_AND_A_NON_LYMPHOID_CELL | Up | 1.46E-41 | 9.82E-39 |
| REACTOME_TRANSLOCATION_OF_ZAP_70_TO_IMMUNOLOGICAL_SYNAPSE | Up | 2.20E-29 | 7.42E-27 |
| **innate lymphocyte** | | | |
| REACTOME_DEADENYLATION_OF_MRNA | Up | 0.00189422 | 0.690751972 |
| REACTOME_ACTIVATION_OF_THE_MRNA_UPON_BINDING_OF_THE_CAP_BINDING_COMPLEX_AND_EIFS_AND_SUBSEQUENT_BINDING_TO_43S | Up | 0.00345795 | 0.690751972 |
| **CD8 GZMK** | | | |
| REACTOME_CHEMOKINE_RECEPTORS_BIND_CHEMOKINES | Up | 4.60E-15 | 3.10E-12 |
| REACTOME_APOBEC3G_MEDIATED_RESISTANCE_TO_HIV1_INFECTION | Up | 1.82E-14 | 6.14E-12 |
| **NK T cells** | | | |
| REACTOME_IMMUNOREGULATORY_INTERACTIONS_BETWEEN_A_LYMPHOID_AND_A_NON_LYMPHOID_CELL | Up | 1.48E-41 | 9.98E-39 |
| REACTOME_OTHER_SEMAPHORIN_INTERACTIONS | Up | 3.88E-12 | 1.31E-09 |
| **NK cells** | | | |
| REACTOME_TRANSLOCATION_OF_ZAP_70_TO_IMMUNOLOGICAL_SYNAPSE | Down | 8.52E-71 | 5.74E-68 |
| REACTOME_PHOSPHORYLATION_OF_CD3_AND_TCR_ZETA_CHAINS | Down | 1.56E-60 | 5.26E-58 |
| **γδ T cells** | | | |
| REACTOME_CHEMOKINE_RECEPTORS_BIND_CHEMOKINES | Up | 2.95E-09 | 1.99E-06 |
| REACTOME_APOBEC3G_MEDIATED_RESISTANCE_TO_HIV1_INFECTION | Up | 1.34E-07 | 4.50E-05 |
| **CD4 Treg** | | | |
| REACTOME_CTLA4_INHIBITORY_SIGNALING | Up | 1.55E-18 | 1.05E-15 |
| REACTOME_IL_RECEPTOR_SHC_SIGNALING | Up | 6.32E-15 | 2.13E-12 |
| **CD4 NFκB** | | | |
| REACTOME_NFKB_IS_ACTIVATED_AND_SIGNALS_SURVIVAL | Up | 1.62E-19 | 1.09E-16 |
| REACTOME_RIP_MEDIATED_NFKB_ACTIVATION_VIA_DAI | Up | 8.19E-14 | 2.76E-11 |
| **CD4 IFN** | | | |
| REACTOME_INTERFERON_ALPHA_BETA_SIGNALING | Up | 0 | 0 |
| REACTOME_INTERFERON_SIGNALING | Up | 1.33E-106 | 2.99E-104 |
| **proliferating NK/T** | | | |
| REACTOME_G1_S_SPECIFIC_TRANSCRIPTION | Up | 1.50E-261 | 1.01E-258 |
| REACTOME_E2F_MEDIATED_REGULATION_OF_DNA_REPLICATION | Up | 3.01E-123 | 1.01E-120 |
| **CD4 TFH** | | | |
| REACTOME_PD1_SIGNALING | Up | 3.17E-36 | 2.14E-33 |
| REACTOME_IL_6_SIGNALING | Up | 2.62E-35 | 8.84E-33 |

*the two most significant (FDR) pathways per functional subcluster are presented

**Supplementary Table E6.** REACTOME analysis of myeloid, B, and epithelial cell clusters

| | Direction | PValue | FDR |
|---|---|---|---|
| **B cells** | | | |
| REACTOME_ANTIGEN_ACTIVATES_B_CELL_RECEPTOR_LEADING_TO_GENERATION_OF_SECOND_MESSENGERS | Up | 1.27E-61 | 8.59E-59 |
| REACTOME_PHOSPHORYLATION_OF_CD3_AND_TCR_ZETA_CHAINS | Up | 4.77E-22 | 1.61E-19 |
| **Basal cells** | | | |
| REACTOME_ENDOSOMAL_VACUOLAR_PATHWAY | Down | 1.13E-185 | 7.58E-183 |
| REACTOME_PHOSPHORYLATION_OF_CD3_AND_TCR_ZETA_CHAINS | Down | 2.79E-156 | 9.40E-154 |
| **cDC2** | | | |
| REACTOME_PHOSPHORYLATION_OF_CD3_AND_TCR_ZETA_CHAINS | Up | 7.89E-118 | 5.32E-115 |
| REACTOME_TRANSLOCATION_OF_ZAP_70_TO_IMMUNOLOGICAL_SYNAPSE | Up | 9.03E-103 | 3.04E-100 |
| **monocytes** | | | |
| REACTOME_CREATION_OF_C4_AND_C2_ACTIVATORS | Up | 3.28E-87 | 2.21E-84 |
| REACTOME_TRANSLOCATION_OF_ZAP_70_TO_IMMUNOLOGICAL_SYNAPSE | Up | 2.19E-79 | 7.39E-77 |
| **goblet/club cell** | | | |
| REACTOME_ENDOSOMAL_VACUOLAR_PATHWAY | Down | 1.92E-165 | 1.30E-162 |
| REACTOME_PHOSPHORYLATION_OF_CD3_AND_TCR_ZETA_CHAINS | Down | 3.95E-146 | 1.33E-143 |
| **enodthelial cell** | | | |
| REACTOME_PHOSPHORYLATION_OF_CD3_AND_TCR_ZETA_CHAINS | Down | 9.90E-103 | 6.68E-100 |
| REACTOME_TRANSLOCATION_OF_ZAP_70_TO_IMMUNOLOGICAL_SYNAPSE | Down | 1.48E-87 | 4.98E-85 |
| **ciliated cell** | | | |
| REACTOME_TERMINATION_OF_O_GLYCAN_BIOSYNTHESIS | Up | 8.55E-04 | 0.440823574 |
| REACTOME_GLUTATHIONE_CONJUGATION | Up | 0.001308082 | 0.440823574 |
| **migratory DC** | | | |
| REACTOME_CHEMOKINE_RECEPTORS_BIND_CHEMOKINES | Up | 7.97E-12 | 2.69E-09 |
| REACTOME_IL_7_SIGNALING | Up | 7.21E-11 | 1.62E-08 |
| **plasmacytoid DC** | | | |
| REACTOME_INTRINSIC_PATHWAY_FOR_APOPTOSIS | Up | 8.74E-26 | 5.89E-23 |
| REACTOME_GLUCAGON_TYPE_LIGAND_RECEPTORS | Up | 4.09E-18 | 1.38E-15 |
| **neutrophil** | | | |
| REACTOME_TRAFFICKING_AND_PROCESSING_OF_ENDOSOMAL_TLR | Up | 4.75E-21 | 3.20E-18 |
| REACTOME_SIGNAL_REGULATORY_PROTEIN_SIRP_FAMILY_INTERACTIONS | Up | 1.40E-16 | 4.72E-14 |
| **cDC1** | | | |
| REACTOME_TRANSLOCATION_OF_ZAP_70_TO_IMMUNOLOGICAL_SYNAPSE | Up | 5.72E-39 | 3.86E-36 |
| REACTOME_PHOSPHORYLATION_OF_CD3_AND_TCR_ZETA_CHAINS | Up | 4.32E-38 | 1.45E-35 |
| **tracheobronchial goblet/club cells** | | | |
| REACTOME_ETHANOL_OXIDATION | Up | 1.04E-18 | 7.02E-16 |
| REACTOME_XENOBIOTICS | Up | 3.32E-13 | 1.12E-10 |
| **dividing B cells** | | | |
| REACTOME_G1_S_SPECIFIC_TRANSCRIPTION | Up | 3.97E-48 | 2.68E-45 |
| REACTOME_E2F_MEDIATED_REGULATION_OF_DNA_REPLICATION | Up | 1.30E-32 | 4.40E-30 |
| **mast cells** | | | |
| REACTOME_NA_CL_DEPENDENT_NEUROTRANSMITTER_TRANSPORTERS | Up | 3.48E-33 | 1.17E-30 |
| REACTOME_DEGRADATION_OF_THE_EXTRACELLULAR_MATRIX | Up | 7.81E-31 | 1.76E-28 |
| **plasma B cells** | | | |
| REACTOME_TRANSLOCATION_OF_ZAP_70_TO_IMMUNOLOGICAL_SYNAPSE | Down | 1.18E-33 | 4.30E-31 |
| REACTOME_SYNTHESIS_SECRETION_AND_DEACYLATION_OF_GHRELIN | Up | 1.34E-33 | 4.30E-31 |

*the two most significant (FDR) pathways per functional subcluster are presented

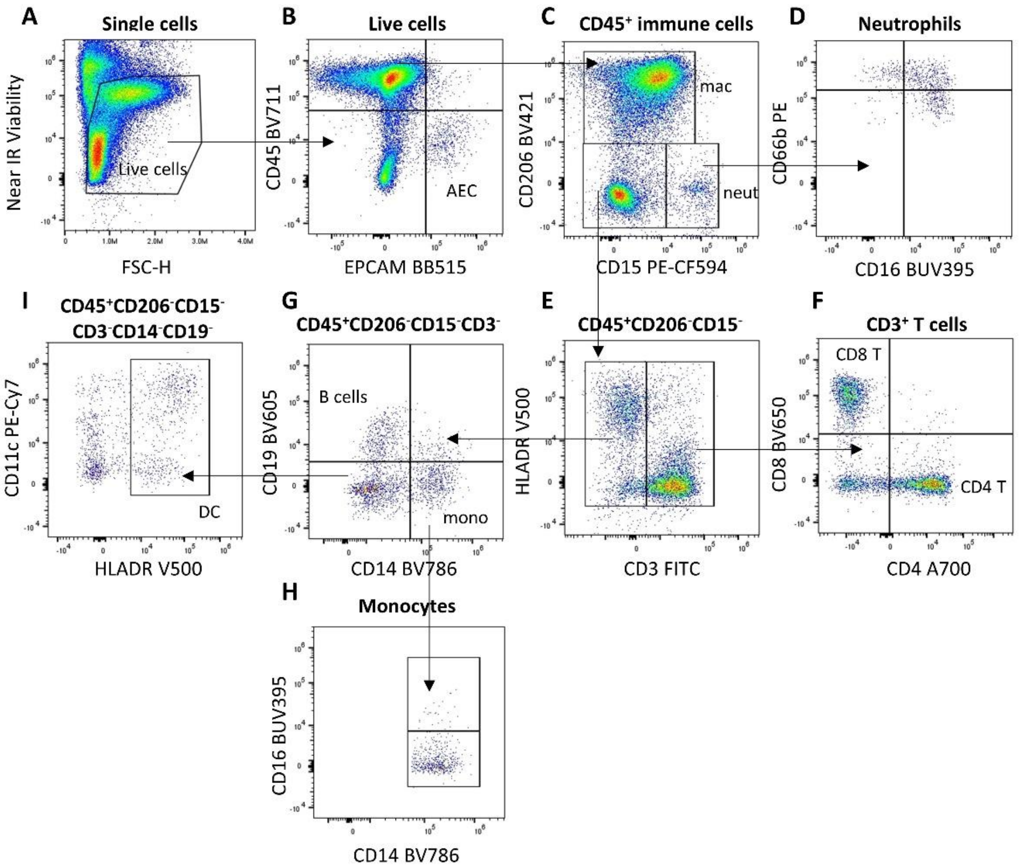


**Supplementary Figure E1.** Representative flow cytometry manual gating of BAL cells. **(A)** Live cells were negative for near IR viability dye. **(B)** Within the live single cell gate, immune cells were $CD45^+$ and airway epithelial cells (AEC) were $CD45^-EPCAM^+$. **(C)** Macrophages were identified within the $CD45^+$ immune cell gate by expression of CD206, and neutrophils were $CD206^-CD15^+$. **(D)** Within the neutrophil populations, three subtypes were identified: $CD16^-CD66b^{high}$, $CD16^+CD66b^{high}$ and $CD16^+CD66b^{low.}$ **(E)** Within the $CD45^+CD206^-CD15^-$ population, T cells were $CD3^+$ and these were then identified as **(F)** $CD4^+$ or $CD8^+$ T cells. **(G)** Within the $CD45^+CD206^-CD15^-CD3^-$ gate, monocytes were identified based on CD14 expression and B cells were $CD14^-CD19^+$. **(H)** A small proportion of monocytes were positive for CD16. **(I)** Finally, cells negative for all preceding markers (excluding CD45) were identified as DCs based on positive staining for HLADR.

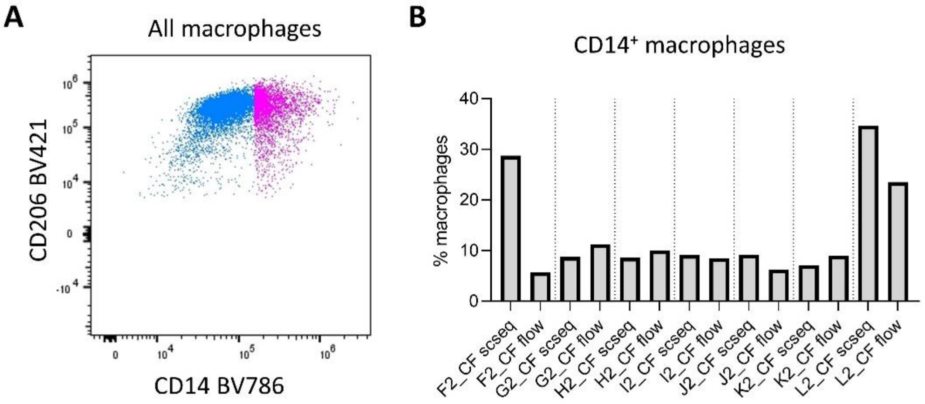


**Supplementary Figure E2.** CD14 expressing macrophages identified by flow cytometry. (A) Manual gating strategy where pink cells represent CD14 expressing macrophages and blue cells represent CD14 negative macrophages. (B) Proportions of $CD14^{+}$ macrophages identified by scRNA-seq and flow cytometry for each individual with both data types collected.
